# Critical assessment of pan-genomics of metagenome-assembled genomes

**DOI:** 10.1101/2022.01.13.476228

**Authors:** Tang Li, Yanbin Yin

## Abstract

**Background:** Large scale metagenome assembly and binning to generate metagenome-assembled genomes (MAGs) has become possible in the past five years. As a result, millions of MAGs have been produced and increasingly included in pan-genomics workflow. However, pan-genome analyses of MAGs may suffer from the known issues with MAGs: fragmentation, incompleteness, and contamination, due to mis-assembly and mis-binning. Here, we conducted a critical assessment of including MAGs in pan-genome analysis, by comparing pan-genome analysis results of complete bacterial genomes and simulated MAGs.

**Results:** We found that incompleteness led to more significant core gene loss than fragmentation. Contamination had little effect on core genome size but had major influence on accessory genomes. The core gene loss remained when using different pan-genome analysis tools and when using a mixture of MAGs and complete genomes. Importantly, the core gene loss was partially alleviated by lowering the core gene threshold and using gene prediction algorithms that consider fragmented genes, but to a less degree when incompleteness was higher than 5%. The core gene loss also led to incorrect pan-genome functional predictions and inaccurate phylogenetic trees.

**Conclusions:** We conclude that lowering core gene threshold and predicting genes in metagenome mode (as Anvi’o does with Prodigal) are necessary in pan-genome analysis of MAGs to alleviate the accuracy loss. Better quality control of MAGs and development of new pan-genome analysis tools specifically designed for MAGs are needed in future studies.

## Background

The term pan-genome was first proposed in bacteria in 2005, representing the entire gene set of all strains in a species [1]. Genes in a pan-genome are classified into three categories: core, accessory, and unique. In theory, core genes must be shared by all strains within the species, accessory genes are present in a subset of the strains, and unique genes are only present in a specific strain [1,2]. In practice, core genes can be defined with a more relaxed threshold. Since 2005, pan-genome analysis has become an essential component of comparative genomics study not only in prokaryotes but also in plants [3], fungi [4], animals [5] and humans [6]. In bacteria, pan-genome analysis has broad applications in studying genomic diversity and phylogeny [7,8], disease outbreak [9], virulence-associated genes [10] and antimicrobial resistance [2,11].

A great number of computational tools for bacterial pan-genome analysis have been developed, such as PGAP [12], GET_HOMOLOGUES [13], ITEP [14], Roary [15], Anvi’o [16], BPGA [17], and PanX [18]. Surveys and comparisons of these tools have been published in recent years [2,11,19,20]. One recent study [21] revealed that the incomplete and inconsistent gene annotations may lead to underestimated core genome size and overestimated pan-genome size. This is particularly important as it is now very common to include draft isolate genomes and metagenome-assembled genomes (MAGs) in pan-genome analysis.

MAGs are produced from metagenome shotgun sequencing reads through filtering, assembling, binning, and taxonomy assignment to generate the approximate representation of actual individual genomes. The term “MAG” first appeared in the literature in 2015 [22,23], when the first large-scale MAG study was published [24]. In 2017, the Genomic Standards Consortium (GSC) published the Minimum Information about a Metagenome-Assembled Genome (MIMAG), a metagenomics community standard for releasing MAGs with mandatory metrics (genome completeness and contamination) [25]. Since then, hundreds of thousands of MAGs have been reconstructed from various environments, e.g., ocean [26], soil [27], freshwater [28], human gut [29,30], activated sludge [31], and animal gut [32,33]. These MAGs are extremely useful to improve predictions of metabolic capacities, discover completely novel taxa, and expand the tree of life, [34,35]. However, a recent study revealed that MAGs with 95% completeness captured only ~77% population core genes and ~50% variable genes, and the quality of MAGs was often worse than expected [36]. Another study [35] critically reviewed concerns that would significantly limit the use of MAGs, such as gaps, assembly and binning errors, chimeras, and contaminations. Even high-quality MAGs (>90% completeness and < 5% contamination according to MIMAG) may still contain assembly errors and chimeras [25,37].

In the past four years, numerous studies have been published using MAGs in pan-genome analysis of human microbiomes [30,38–41]. Obviously, it has become a routine to combine MAGs with isolate genomes or use only MAGs for pan-genome analysis. However, to which extent the accuracy of pan-genome results may be influenced by the caveats (fragmentation, incompleteness, and contamination) of MAGs has never been critically assessed. We hypothesized that including MAGs will lead to biases and errors in pan-genome analysis results. We tested this hypothesis by comparing pan-genome analysis results of complete genomes and MAGs simulated from complete genomes by introducing fragmentation, incompleteness, and contamination. Based on our findings, we have provided recommendations on how to alleviate the accuracy loss due to the use of MAGs.

## Results

### MAGs are increasingly used in pan-genome analyses in various ecological environments

Since 2017, there was an increasing use of MAGs in pan-genome analyses (**Figure 1A**). This number is certainly underestimated as our literature search applied very strict conditions (see **Methods**). These analyzed MAGs were from various ecological environments (**Figure 1B**). The most popular pan-genome analysis tools were Anvi’o [16] and Roary [15] (**Figure 1C**), while other tools such as BPGA [17], GET_HOMOLOGUES [13], and OrthoMCL [42] were also used. The strictest core gene (CG) threshold 100% (see **Methods**) was used in 2/3 studies (**Figure 1D**). In some studies, only MAGs were used in the pan-genome analyses (MAG% = 100% in **Figure 1E**), e.g., the studies of hydrothermal vents [43,44], ocean [45], and animal gut [46]. However, in others a mixture of complete isolate genome and MAGs were used (MAG% < 100% **Figure 1E**), e.g., in human gut [40,47]. Some studies analyzed pan-genomes of thousands of genomes/MAGs (**Figure 1F**), while most studies analyzed < 100 genomes/MAGs. The details about these publications are provided in **Table S1**.

**Figure 1.**
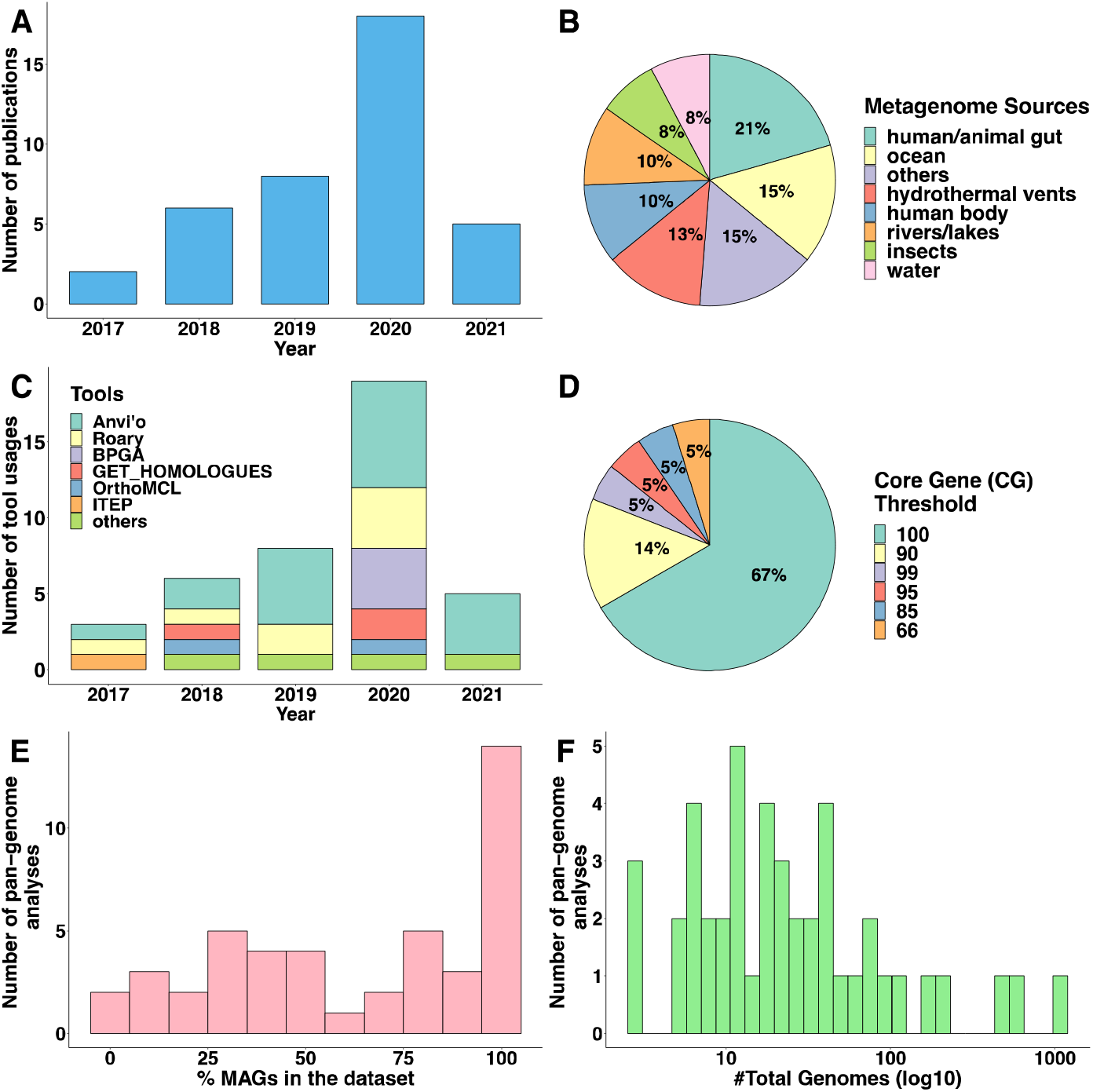
Summary of 39 publications that used MAGs in pan-genome analysis. (**A**): The bar plot shows the publication years. (**B**): The pie chart shows the ecological environments of analyzed MAGs. (**C**): The bar plot shows the pan-genome analysis tools (multiple tools used in a same study were counted separately). (**D**): The pie chart shows the CG thresholds used in pan-genome studies. (**E**): The bar plot to survey the percentages of MAGs used in pan-genome analysis (y-axis is the count of pan-genomes, and x-axis is the percentages of MAGs). (**F**): The bar plot to survey the numbers of genomes/MAGs used in pan-genome analysis (y-axis is the count of pan-genomes, and x-axis is the numbers of genomes/MAGs).

### Pan-genome analysis of complete genomes of 17 bacterial species finds a positive correlation between pan-genome size and the average genome size

The 17 species used in this study belong to 15 taxonomic families of three phyla: Proteobacteria (10 species), Firmicutes (5) and Actinobacteria (2) (**Table 1)**. All the complete genomes were listed in **Table S2**. The pan-genome size (**Table 1**, **Figure S1**) was positively correlated with the average genome size (Pearson correlation coefficient R = 0.72, p = 0.0012) [21,48] and the number of genomes used (R = 0.71, p = 0.0013). One well-known fact about pan-genome analysis is that some bacterial species has more open pan-genome than others [49,50]. The Heaps’ law model was often used to predict the openness and closeness of the pan-genome [49,51]. The γ values shown in **Table 1** are consistent with previous findings [52–54]: the two largest γ values were found in *S. enterica* and *E. coli*, while three species (*B. pertussis*, *M. tuberculosis*, and *C. pseudotuberculosis*) only had γ values at ~0.03. The lower γ values indicate that their pan-genomes were near to be closed. Since *E. coli* and *B. pertussis* not only have larger numbers of complete genomes but also have extreme differences in ANI values, pan-genome structures, and γ values, they were selected as representative species in this study.

**Table 1.** Summary of 17 bacterial species.

| # | Species | Number of Complete Genomes in RefSeq | Family | Phylum | Genome Size (Average # Genes) | ANI (%) | Pan-genome Size<sup>a</sup> (# Gene Families) | Core Genome Size<sup>a</sup> (# Gene Families) | $\gamma$ value<sup>b</sup> |
| --- | --- | --- | --- | --- | --- | --- | --- | --- | --- |
| 1 | *Escherichia coli* | 923 | Enterobacteriaceae | Proteobacteria | 4873 | 97.78 | 33123 | 1486 | 0.3170 |
| 2 | *Salmonella enterica* | 746 | Enterobacteriaceae | Proteobacteria | 4577 | 98.38 | 27085 | 1877 | 0.3182 |
| 3 | *Bordetella pertussis* | 539 | Alcaligenaceae | Proteobacteria | 3901 | 99.98 | 4115 | 2960 | 0.0357 |
| 4 | *Staphylococcus aureus* | 443 | Staphylococcaceae | Firmicutes | 2639 | 98.43 | 6177 | 1710 | 0.1503 |
| 5 | *Klebsiella pneumoniae* | 372 | Enterobacteriaceae | Proteobacteria | 5320 | 99.04 | 22106 | 2741 | 0.2739 |
| 6 | *Pseudomonas aeruginosa* | 196 | Pseudomonadaceae | Proteobacteria | 6175 | 98.90 | 19637 | 3734 | 0.2669 |
| 7 | *Streptococcus pyogenes* | 195 | Streptococcaceae | Firmicutes | 1758 | 98.77 | 3959 | 1267 | 0.1667 |
| 8 | *Listeria monocytogenes* | 180 | Listeriaceae | Firmicutes | 2909 | 96.96 | 6109 | 2098 | 0.1515 |
| 9 | *Mycobacterium tuberculosis* | 176 | Mycobacteriaceae | Actinobacteria | 4067 | 99.92 | 4449 | 3429 | 0.0316 |
| 10 | *Campylobacter jejuni* | 171 | Campylobacteraceae | Proteobacteria | 1746 | 97.81 | 4027 | 1204 | 0.1865 |
| 11 | *Helicobacter pylori* | 166 | Helicobacteraceae | Proteobacteria | 1545 | 95.25 | 3755 | 1143 | 0.2192 |
| 12 | *Acinetobacter baumannii* | 161 | Moraxellaceae | Proteobacteria | 3813 | 98.28 | 12426 | 2132 | 0.2649 |
| 13 | *Enterococcus faecium* | 125 | Enterococcaceae | Firmicutes | 2937 | 98.39 | 7652 | 1608 | 0.2226 |
| 14 | *Neisseria meningitidis* | 106 | Neisseriaceae | Proteobacteria | 2042 | 98.20 | 3347 | 1414 | 0.1285 |
| 15 | *Bacillus subtilis* | 102 | Bacillaceae | Firmicutes | 4178 | 98.44 | 8987 | 3063 | 0.1901 |
| 16 | *Burkholderia pseudomallei* | 101 | Burkholderiaceae | Proteobacteria | 5918 | 99.41 | 12941 | 4663 | 0.2072 |
| 17 | *Corynebacterium pseudotuberculosis* | 100 | Corynebacteriaceae | Actinobacteria | 2127 | 99.35 | 2449 | 1887 | 0.0370 |

<sup>a</sup>Pan-genome size and core genome size were calculated by using Roary (-i 90 -cd 100 -s -e -n). <sup>b</sup>The $\gamma$ values were calculated by using Heaps' law model [51].

### Fragmentation and incompleteness of MAGs lead to significant core gene loss

To guide the creation of simulated MAGs from complete genomes, we have plotted the distributions of contig number, genome completeness, and contamination in 276,349 UHGG (Unified Human Gastrointestinal Genome) MAGs (**Figure S2A)** [38]. The number of contigs in these real MAGs varied from 1 to 2282 with a mean = 208. The completeness ranged between 50% and 100% with a mean = 85.18%, while the contamination rate varied from 0% to 5% with a mean = 1.2%. These three metrics all have skewed distributions, even when they were plotted for individual species (**Figure S2B**). Therefore, the F-distribution, a theoretical distribution in Statistics used to model skewed distributions, was applied to guide the creation of simulated MAGs from complete genomes (see **Methods** and **Figure 2A**). Overall, for each species, four types of datasets were generated: (1) original datasets (100 complete genomes), (2) fragmentation datasets (fragmented MAGs), (3) incompleteness datasets (fragmented+incomplete MAGs), and (4) contamination datasets (fragmented+incomplete+contaminated MAGs).

**Figure 2.**
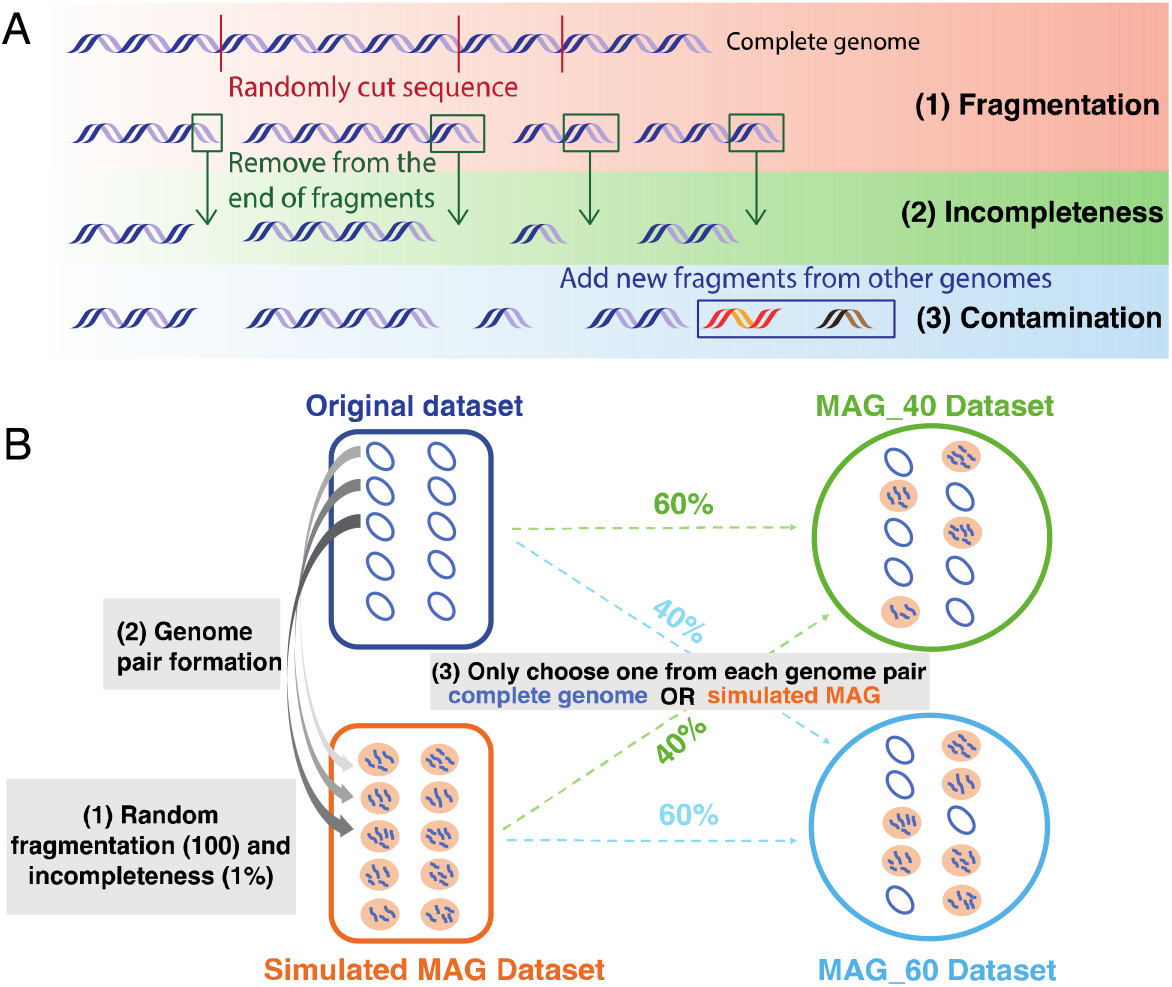
Workflows to generate simulated MAG datasets. (**A**): The pipeline to generate fragmented, incomplete and contaminated MAGs from complete genomes. The first step fragmentation can be performed at 5 different levels (50, 100, 200, 300, and 400 fragments). The second step incompleteness can be performed at 5 different levels (1, 2, 3, 4, and 5%). The third step contamination can be performed at 8 different levels (0.5, 1, 1.5, 2, 2.5, 3, 3.5, 4%). (**B**): Combining complete genomes and simulated MAGs to form mixed datasets. Shown in the diagram as an example, the original dataset contains 100 randomly selected complete genomes. Genomes in the simulated MAG dataset are generated from complete genomes by fragmentation and incompleteness simulation (described in A). Each genome pair contains one complete genome and its corresponding MAG. Two mixed datasets are shown as examples. MAG_40 is created by randomly combining complete genomes (60%) in the original dataset and MAGs (40%) from the simulated MAG dataset. If a complete genome has been chosen, its corresponding MAG would not be selected.

To study the effect of fragmentation and incompleteness, Roary was run with 90% sequence identity and 100% core gene thresholds on the simulated MAG datasets (fragmentation dataset and incompleteness dataset) as well as the original dataset. As expected, the number of core gene families decreased with more fragmented MAGs in *E. coli* (**Figure 3A**) and *B. pertussis* (**Figure 3B**). The reduction of core genome sizes was also observed in other 15 species (**Figure S3A**). To quantitatively evaluate the degree of core gene loss in different species, an exponential model was used (see **Methods** and **Figure 3**). In general, species with a larger number of core gene families (e.g., *B. pseudomallei*) in the original genomes tend to have more rapid core genome reduction.

**Figure 3.**
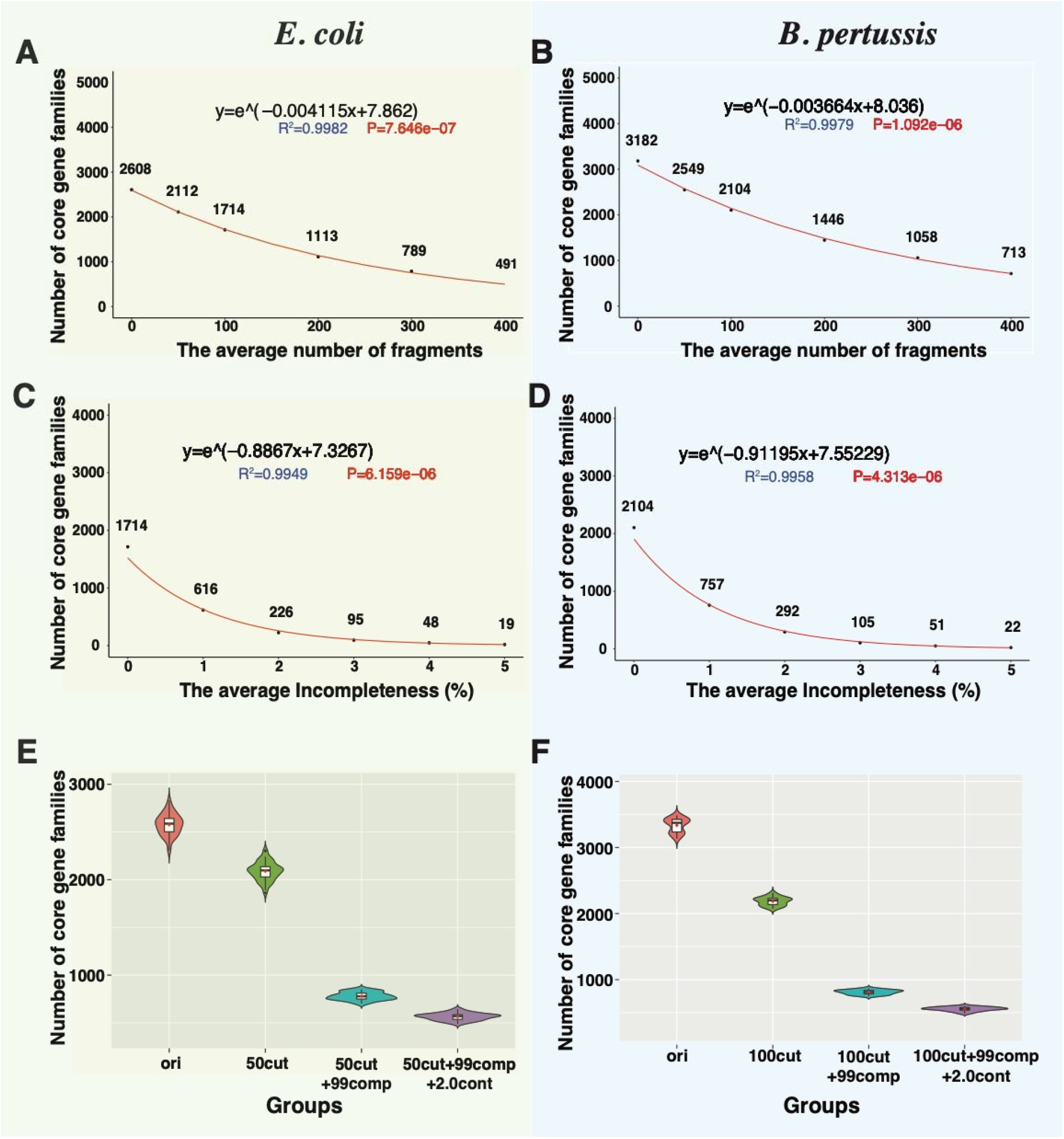
Core genome sizes decrease in simulated MAGs of *Escherichia coli* and *Bordetella pertussis*. (**A**) and (**B**): The core genome size continuously decreases as the simulated MAGs become more fragmented. The red curve was predicted using an exponential model for the correlation between the x-axis (the number of fragments) and the y-axis (the core genome size). (**C**) and (**D**): The core genome size decreases more rapidly as the simulated MAGs become less complete. (**E**): The violin plot of the core genome sizes in 50 *E. coli* original datasets and their corresponding simulated MAG datasets. (**F**): The violin plot of the core genome sizes in 30 *B. pertussis* original datasets and their corresponding simulated MAG datasets. Groups: “ori” represents the original datasets; “50cut” or “100cut” represents the fragmentation datasets; “50cut+99comp” or “100cut+99comp” means that genomes in a dataset have an average of 1% incompleteness based on 50 or 100 fragmentation; “50cut+99comp+2.0cont” or “100cut+99comp+2.0cont” means that genomes in a dataset have an average of 2.0% intra-species contamination based on 50 or 100 fragmentation and 1% incompleteness. All the core genome sizes are calculated by using Roary with 90% identity and 100% core gene thresholds.

Compared to fragmentation, incompleteness (incomplete MAGs with a fixed fragmentation rate: average 100 contigs in each MAG) had more significant effects on the core genome size. Only 36% (616/1,714 in *E. coli* and 757/2,104 in *B. pertussis*) of core gene families were retained in these two species when an average of 1% genome sequences was removed (**Figure 3C, 3D**). Similar results were observed in all other species regardless of their original core genome sizes (**Figure S3B**). Overall, a 1% loss in completeness, from 100% to 99%, would lead to > 60% loss of core genes in most species. A 5% loss in genome completeness (**Figure 3C, 3D, S3B**) would almost lose all core genes.

Choosing which 100 complete genomes to use as the original dataset to generate simulated MAGs may affect the pan-genome analysis results. To assess this effect, we created 50 original datasets for *E. coli*, each dataset with 100 randomly selected complete genomes. For each of the 50 original datasets, we further generated the fragmentation dataset (fragmented MAGs), the incompleteness dataset (fragmented+incomplete MAGs), and the contamination dataset (fragmented+incomplete+contaminated MAGs) (see **Methods**). Overall, the decrease of core gene families in simulated MAGs was observed in every simulated dataset in *E. coli* (**Figure 3E**), *B. pertussis* (**Figure 3F**), and in other species (**Figure S4**).

### Contamination has little effect on core genomes but influences the accessory genomes

Real MAGs are built by binning metagenome contigs based on similar sequence compositions and sequencing coverage [35]. Therefore, contamination is most likely from closely related genomes (e.g., strains of the same species or genus) as they share more similar sequence compositions. As mentioned in **Methods**, we simulated contaminated MAGs by adding sequence fragments from the same species or from different species of the same genus, i.e., intra- or inter-species contaminations (**Figure 2A**). Unlike fragmentation and incompleteness, which caused uniform core gene loss in all species, intra-species contamination led to different core gene changes in different species (**Figure S5A**). In about half species like *B. pseudomallei* and *K. pneumoniae*, a 4% contamination caused ~500 core gene loss. However, there were no noticeable changes in *H. pylori* and *S. pyogenes*.

Intuitively, contamination will result in a larger pan-genome as additional genes are added, but should not reduce the core gene number as no core genes are removed. The unexpected core gene loss is further explained in the next section. However, as expected, in most species, the number of cloud genes (genes that are shared by <15% genomes, defined by Roary) increased constantly when more intra-species contamination was introduced (**Figure S5A**).

Furthermore, when inter-species contamination was introduced, the number of core genes dropped less significantly, whereas the number of cloud genes increased more significantly (**Figure S5B**).

### Core gene loss remains when using different pan-genome analysis tools and mixed MAG datasets

To determine whether the core gene loss was caused by the use of Roary, all the analyses were repeated by using two other popular tools: BPGA [17] and Anvi’o [16] on 10 *E. coli* and 10 *B. pertussis* datasets (i.e., 10 original datasets and their corresponding simulated MAG datasets) with the same gene models that were used by Roary (i.e., predicted by Prokka [55]). The core gene loss remained for fragmentation and incompleteness, but not for contamination (**Figure 4A, 4B**). In fact, in both *E. coli* and *B. pertussis*, with intra-species contamination the number of core genes increased slightly when using BPGA and Anvi’o, as we expected (see above). We hypothesized that the opposite result by Roary (**Figure 3E and 3F**) was caused by possible bias in the gene clustering step of Roary. To test it, the core gene sets before and after a 2% contamination were manually compared. The result indicated that some genes identified as core genes before contamination were misclassified as soft-core genes (genes present in > 95% but < 100% of the genomes) after contamination. In other words, in Roary core genes in some genomes were clustered with non-core genes (e.g., from contamination sources) that share higher sequence identity to form a new gene family, leading to the loss of core gene families.

**Figure 4.**
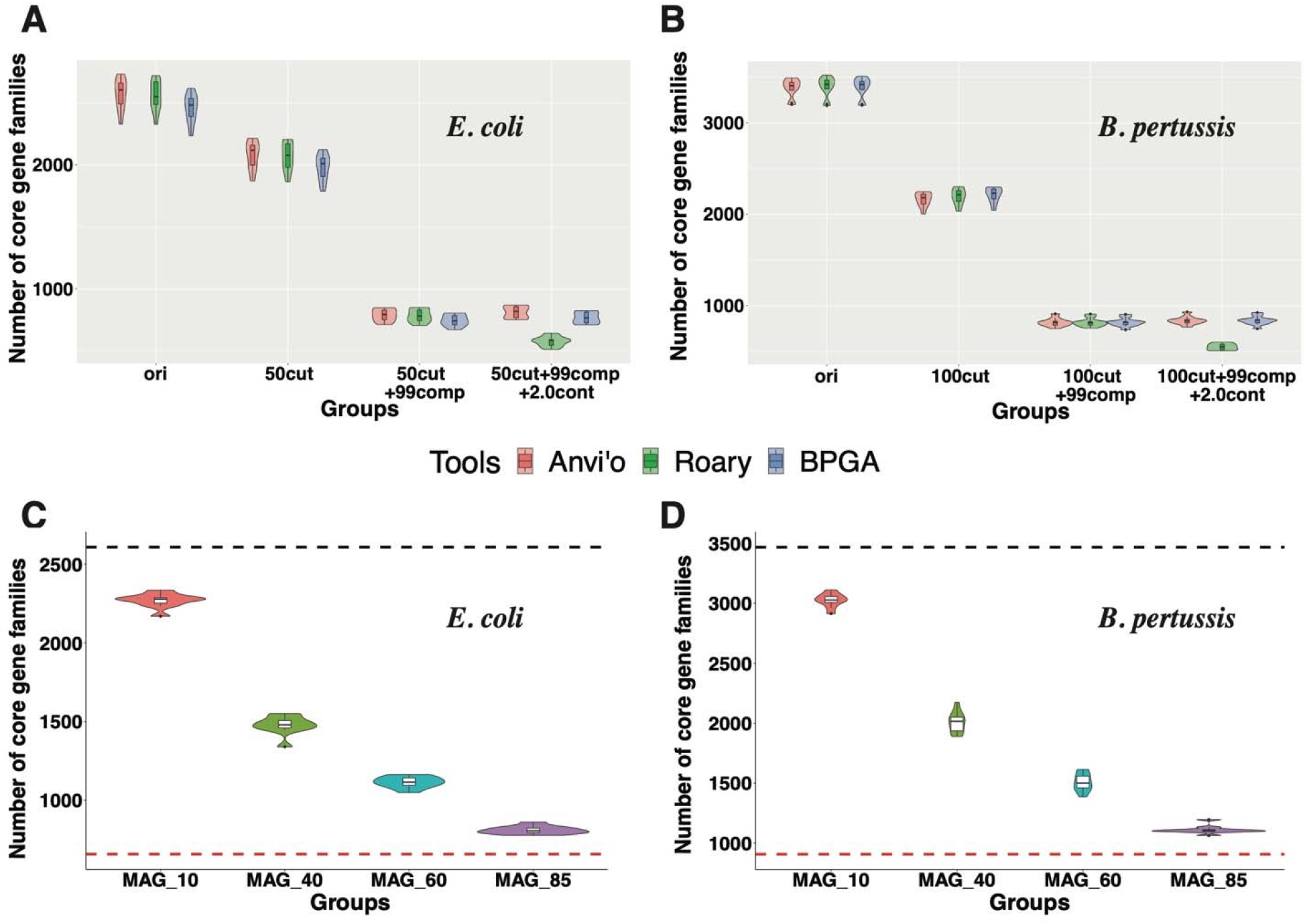
The core genome size change using different pan-genome tools and using mixed MAG datasets. (**A**) and (**B**): The violin plots of the number of core gene families were calculated by Anvi’o (red), Roary (green), and BPGA (blue) in 10 *E. coli* datasets and 10 *B. pertussis* datasets. See more details in legend in **Figure 3E and 3F**. (**C**) and (**D**): The number of core gene families in *E. coli* and *B. pertussis* using mixed MAG dataset (see Methods). MAG_10 means that datasets contain 10% simulated MAGs and 90% complete genomes (see **Figure 2B** for examples). The black dot line indicates the number of core gene families in the original dataset (100% complete genomes). The red dot line indicates the number of core gene families in the simulation dataset (100% MAGs). All the core genomes were predicted by Roary, and each group contains 10 datasets.

As shown in **Figure 1E**, a large proportion of published studies used mixed datasets (mixture of complete genomes and MAGs) in pan-genome analyses. To determine how much core gene loss the mixed datasets may cause, four groups of mixed datasets containing from 10% to 85% MAGs were used in Roary analyses. We found that the higher MAG percentage is included, the more core genes will be lost (**Figure 4C**,**4D**), which fits our expectation.

### Core gene loss can be partially alleviated by lowering the core gene threshold

The core gene (CG) threshold and the sequence identity (SI) are two important parameters in pan-genome analysis (see **Methods**). In above analyses, the CG threshold was set at 100% and the SI threshold was set at 90%. To determine the two parameters’ impacts on pan-genome results, different CG thresholds and different SI thresholds were tested. First, the SI threshold was fixed at >=90%, and only the CG threshold was altered. Under a specific fragmentation level or incompleteness level, a lower CG threshold always gave a higher core gene number (**Figure 5**). This means the core gene loss can be alleviated by the lowering the CG threshold. However, the degree of the alleviation varied depending on what tools were used and how fragmented and incomplete the MAGs were.

**Figure 5.**
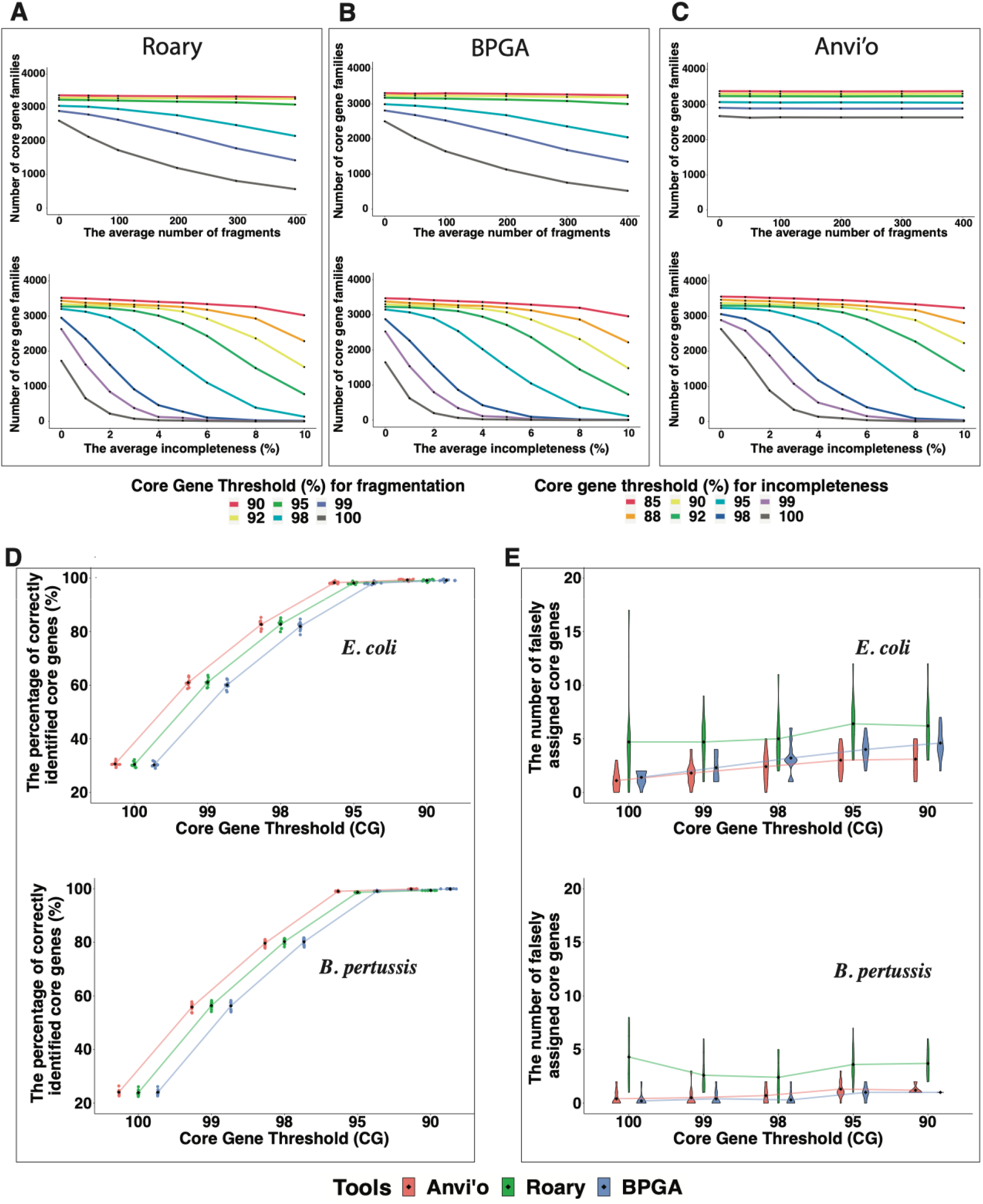
Lowering core gene thresholds help alleviate the core gene loss. **(A-C):** The numbers of core gene families in simulated MAGs of *E. coli* (with different fragmentation and incompleteness levels) were predicted by using Roary, BPGA, and Anvi’o using different core gene thresholds. (**D**): The sinaplots show the impact of lowering CG thresholds in recovering true positives. (**E**): The violin plots show the impact of lowering CG thresholds in introducing false positives. Ten *E. coli* and *B. pertussis* simulated MAG datasets with an average of 50/100 fragmentation and 1% incompleteness were used.

In Roary (**Figure 5A**) and BPGA (**Figure 5B**), a continuous and significant core gene loss was observed in more fragmented MAGs when using CG threshold >= 98%. In contrast, in Anvi’o with its default gene prediction method, more fragmented MAGs did not result in more core gene loss, although choosing lower CG thresholds also resulted in higher core gene numbers (**Figure 5C**). This is because by default Anvi’o uses Prodigal [56] in its metagenome mode for gene prediction, whereas BPGA and Roary use Prodigal (integrated in Prokka) in normal mode for gene prediction. The metagenome mode of Prodigal works for fragmented genes while normal mode does not. Therefore, in Anvi’o, fragmentation had little effect on core gene loss.

Compared to fragmentation, incompleteness caused a more significant core gene loss in all the three tools. Even when the core gene threshold is as low as 85%, for MAGs with an average incompleteness >10%, the core gene loss was still a big problem for all the three tools (**Figure 5A,B,C**). Anvi’o was less affected (**Figure 5C**) but still suffers when incompleteness was >=5% and CG threshold was >=95%. Clearly a higher incompleteness will significantly reduce the size of the core genome regardless of what tools are used.

When using a lower CG threshold, some previous soft-core genes may now become core genes in both original complete genomes and simulated MAGs. Under the same CG threshold, a small number of core genes (<10) were found in only simulated MAGs but not in original complete genomes, and thus are false positives; core genes that were found in both complete genomes and simulated MAGs are true positives. With the CG threshold decreasing, the false positive rate rose only slightly (**Figure 5E**), but the true positive rate rose very significantly (**Figure 5D**), almost to 100% when the CG threshold was lowered to <=95% (100 fragments and 1% incompleteness). Therefore, using a lower CG threshold based on the quality of MAGs helped recover the lost core genes with very low false positive rate.

In contrast, changing the SI threshold (with fixed CG threshold) had very little help in recovering lost core genes in simulated MAGs (**Figure S6)**.

### Core gene loss leads to underestimation in core gene functional analysis

In pan-genome analysis, after the core and accessory genes are identified, downstream analyses are often performed on these genes to better understand the evolution of different genomes and functions of different genes. In addition, functional enrichment analysis is also often performed on core genes, accessory genes, and unique genes to explain species adaptation to various environments. Given that the core genome size is inevitably decreased in the pan-genome analysis of MAGs, there will be some consequences in the COG functional predictions for the core genes. When using the 100% CG threshold, the number of core genes assigned to COG categories was dramatically decreased with increasing fragmentation and incompleteness (**Figure 6**). Incompleteness (**Figure 6C, 6D**) had a more significant impact than fragmentation (**Figure 6A, 6B**). More than 50% of the core gene representatives were lost in each COG category with only 1% incompleteness (**Figure 6C, 6D**). When using the 100% CG threshold, the COG functional enrichment in the core and accessory genes were also significantly changed due to fragmentation and incompleteness (**Figure S7A-C**), leading to inaccurate inferences to explain environmental adaptation.

**Figure 6.**
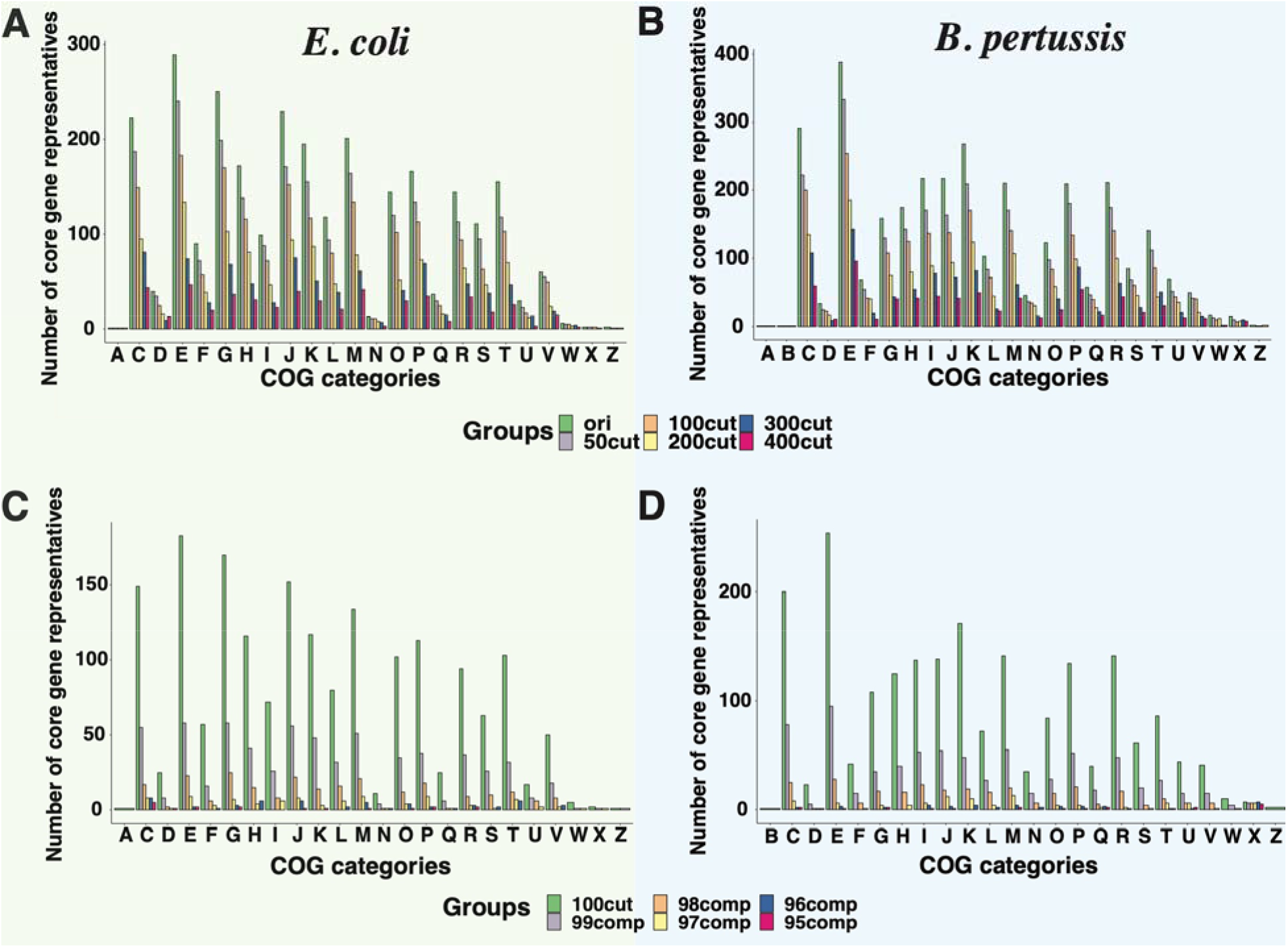
The COG analysis of core genes in simulated MAGs. (**A**) and (**B**): The bar plots of the number of core genes with different fragmentation levels in each COG category. (**C**) and (**D**): The bar plots of the number of core genes with different incompleteness levels in each COG category. COG category single letter codes: A: RNA processing and modification, C: Energy production and conversion, D: Cell cycle control and mitosis, E: Amino Acid metabolism and transport, F: Nucleotide metabolism and transport, G: Carbohydrate metabolism and transport, H: Coenzyme metabolism, I: Lipid metabolism, J: Translation, K: Transcription, L: Replication and repair, M: Cell wall/membrane/envelop biogenesis, N: Cell motility, O: Post-translational modification, protein turnover, chaperone functions, P: Inorganic ion transport and metabolism, Q: Secondary Structure, R: General Functional Prediction only, S: Function Unknown, T: Signal Transduction; U: Intracellular, trafficking and secretion, V: Defense mechanisms, W: Extracellular structures, X: Mobilome: prophages, transposons, Z: Cytoskeleton.

As expected, when using lower core gene thresholds, less significant decreases were observed on core gene functional predictions caused by fragmentation and incompleteness (**Figure S8).** Moreover, the COG functional enrichment analysis of core and accessory genes were also less affected (**Figure S7D-F**). However, the accuracy loss could not be fully eliminated especially for MAGs with higher incompleteness. It is also likely that the inclusion of falsely predicted core genes may lead to incorrect functional analysis results.

### Simulated MAGs cause significant change in phylogenetic tree topology

Pan-genome analysis tools often produce a gene presence and absence matrix, which has rows representing all the genomes and columns representing the different gene clusters (i.e., gene families). From this matrix, a phylogenetic tree can be constructed to delineate the evolutionary relationship among the studied genomes. When the genomes were fragmented, incomplete and contaminated, the gene presence and absence matrix will change, and then the phylogenetic tree will be affected. Indeed, comparing to the complete *E. coli* genomes (**Figure 7A**), a dramatic shrinkage in the core gene area (dark blue area shown in the red frame) was observed in the gene presence and absence dot plot of simulated *E. coli* MAGs (**Figure 7B**). The overall tree topology also changed, although some clades (e.g., the blue and purple branches in **Figure 7A, 7B**) were more conserved than others.

**Figure 7.**
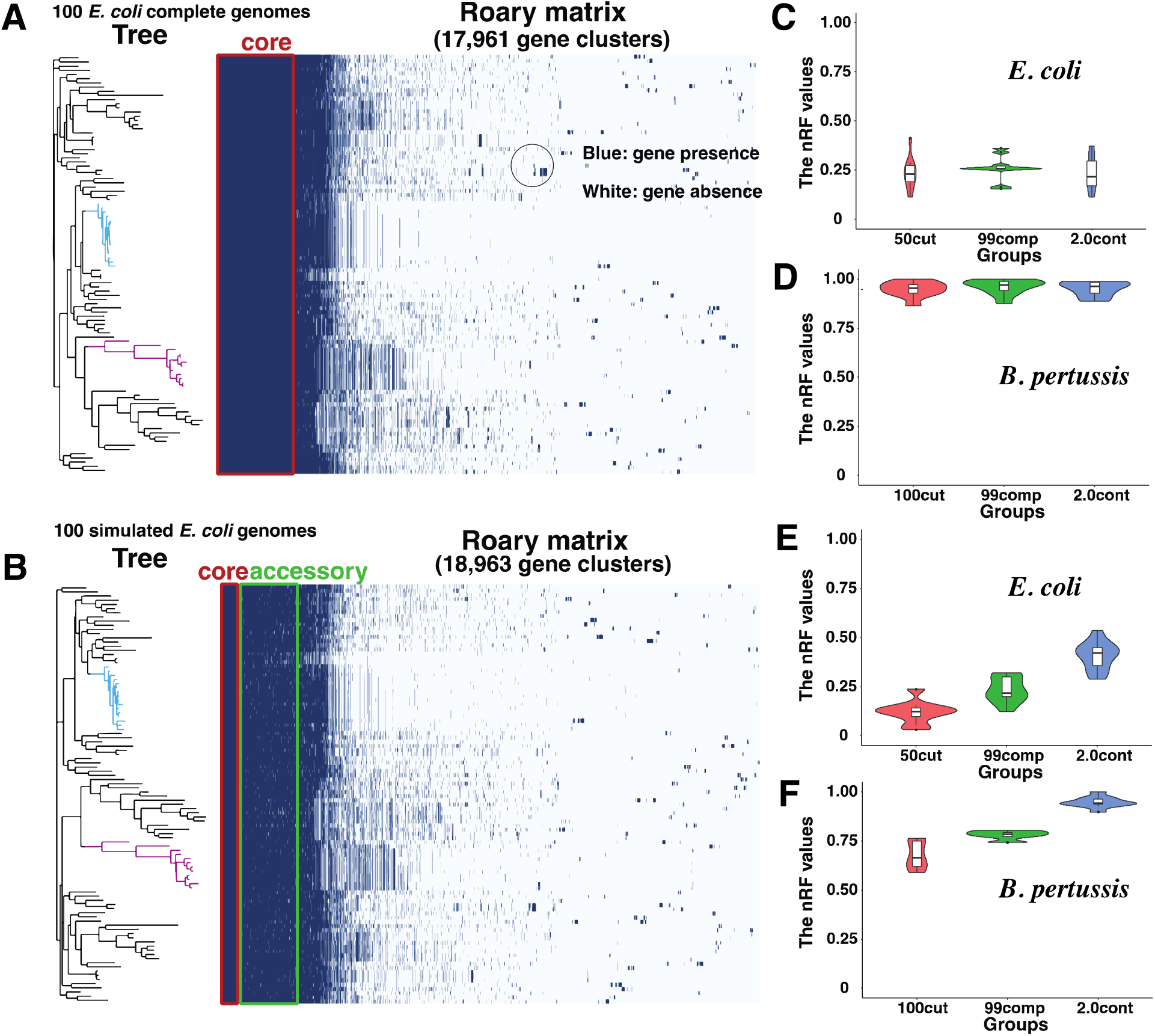
Simulated MAGs cause significant change in phylogenetic tree. (**A**): The phylogenetic tree and Roary gene presence and absence matrix (as a dot plot) for 100 *E. coli* complete genomes. (**B**): The phylogenetic tree and Roary gene presence and absence matrix (as a dot plot) for 100 *E. coli* simulated MAGs with an average of 100 fragments and 1% incompleteness. In the dot plot, each row corresponds to a branch (one genome or MAG) in the tree, and each column represents a gene family/cluster. White indicates gene absence and blue indicates gene absence. (**C**) and (**D**): The violin plots of the nRF distance between MAG tree and complete genome tree constructed based on gene presence and absence matrix (10 *E. coli* datasets). (**E**) and (**F**): The violin plots of the nRF distance values between MAG tree and complete genome tree constructed based on core genome alignment (10 *E. coli* datasets).

To quantify the topological changes between the trees before and after simulation, the normalized Robinson-Foulds (nRF) distance was used. The higher the nRF values, the more differences between the trees. The nRF distance values varied between 0.1 and 0.4 among 10 *E. coli* datasets in different simulation groups (**Figure 7C**); however, these values were larger than 0.85 in *B. pertussis* datasets (**Figure 7D**). Similarly, low nRF values were observed in *S. aureus* (**Figure S9A**) while high nRF values were found in *B. pseudomallei* (**Figure S9B**). These suggest that different species are affected by MAGs to a different degree. In addition to the gene presence and absence matrix, phylogenetic trees can also be built based on core gene alignment. The nRF values increased from 0.1 in simulated MAGs with only fragmentation to 0.4 in simulated MAGs with fragmentation+incompleteness+contamination in *E. coli* datasets (**Figure 7E**), whereas these values changed from 0.6 to 0.9 in *B. pertussis* datasets (**Figure 7F**). In contrast, the percentage of branches shared between trees in simulation and original groups significantly decreased (**Figure S9C, S9D**). All these indicate that the more fragmentation, incompleteness and contamination MAGs have, the less accurate the phylogeny will be, irrespective of using gene presence and absence or core gene alignment for tree construction.

## Discussion

Concerns have been recently raised that mis-assembly and mis-binning can be very common but difficult to identify in the automatically generated MAGs [25,36,37,57,58]. Given that MAGs are known to be fragmented, incomplete, and in many cases also contaminated, we hypothesized that pan-genomics may be significantly affected by including MAGs. In this study, we aimed to critically evaluate the accuracy loss of using MAGs in pan-genomics. We approached this goal by comparing the pan-genome analysis results of complete genomes and simulated MAGs. Our simulation considers fragmentation, incompleteness, and contamination (**Figure 2**), and follows the empirical distribution (**Figure S2**) of real MAGs from human gut microbiome [38].

Our major finding is that the core gene loss caused by fragmentation and incompleteness in MAGs is universal, regardless of what species is used (**Figure 3** and **Figure S3**), what pan-genome analysis software is used (**Figure 4**), what core gene threshold is used (**Figure 5**), and what fraction of MAGs is included (**Figure 4**). However, to what extent the core gene can be lost in MAGs varies among species, softwares, core gene thresholds, and fractions of MAGs. The most important finding is that lowering the core gene threshold and choosing the metagenome mode for gene prediction (as Anvi’o does) can alleviate the core gene loss (**Figure 5A,B,C**) with minimal false positives (**Figure 5D,E**).

Why fragmentation, incompleteness, and even contamination can cause core gene loss? Previous studies have shown that pan-genome analysis suffers from incomplete, inconsistent, and inaccurate gene predictions in bacterial isolate genomes [21,59]. Therefore, the core gene loss in MAGs must be directly caused by missing genes in the gene prediction step. Shown in **Figure 8**, we delineated the process of core gene loss with fragmentation, incompleteness, and contamination. Firstly, if the fragmentation disrupts a gene coding region, the gene will be lost due to the missing of start or stop codons. Secondly, more genes might be partially or fully removed due to deletion caused by incompleteness. For both processes, gene prediction programs will fail to call the affected genes, or call an incorrect gene due to reading frame shifts, leading to falsely predicted genes [60]. Thirdly, contamination from closely related strains or species should not lead to core gene loss due to no removal of existing genomic regions. However, the size of the core genome still decreased when using Roary, which was not seen in BPGA and Anvi’o (**Figure 4A,B**). It appears that some core gene clusters (families) before contamination were falsely split into multiple gene clusters in Roary, leading to the loss of core genes. Gene clusters being incorrectly split into multiple smaller clusters were also noticed in other studies [21,59], which suggested that the removal of contamination and annotation errors are essential to construct an accurate pan-genome [59]. Compared to these previous works, our study went further and generated simulated MAGs resembling real MAGs and also considered fragmentation and incompleteness.

**Figure 8.**
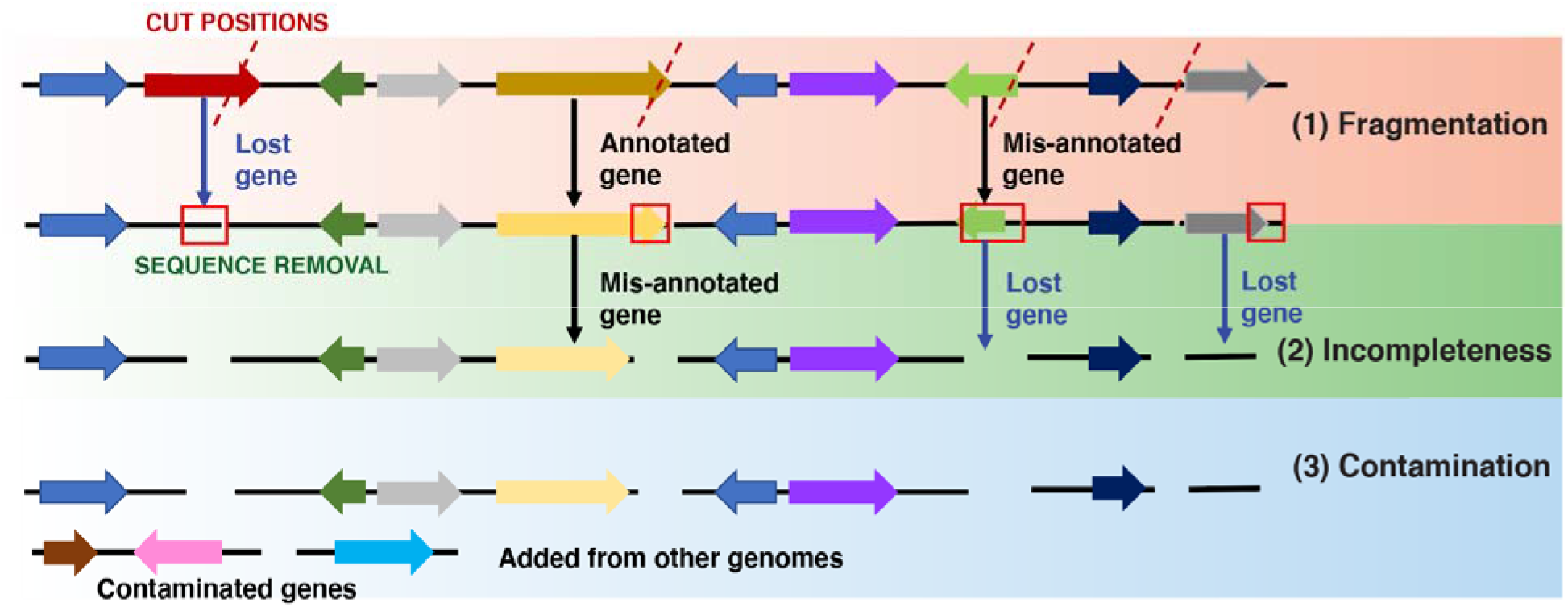
Incorrect gene predictions caused by lost genes or falsely predicted genes due to fragmentation, incompleteness, and contamination. Arrows shown in different colors represent predicted genes in genome sequences.

What is the solution to core gene loss in pan-genomics of MAGs? First, the strictest core gene threshold (i.e., genes present in 100% genomes) should not be used for MAGs. According to **Figure 5**, a higher CG threshold like 95% can be applied to MAGs with only low fragmentation and low incompleteness. However, a lower CG threshold like 90% or 85% should be used for MAGs with higher fragmentation (average number of fragments > 100), and higher incompleteness (average incompleteness > 5%). Although the clustering sequence identity is also an important parameter in pan-genome analysis, changing the threshold of it did not affect the core gene counts in MAGs (**Figure S6**). The percentage of MAGs used in mixed datasets is another important factor. Mixing MAG with complete isolate genomes will reduce the core genome loss (**Figure 5**). Second, it is important to use gene prediction programs that can handle fragmented genes. Some pan-genome analysis tools call gene prediction programs internally (e.g., Anvi’o), while others demand users to provide externally predicted genes as input (e.g., Roary and BPGA). This study primarily used Prokka to prepare predicted genes as input to Anvi’o, Roary and BPGA. However, when Anvi’o was run with its default gene prediction method (metagenome mode of Prodigal), the core gene loss due to increasing fragmentation can be almost negligible (**Figure 5**). Third, some very recent pan-genome analysis tools, e.g., Panaroo [59] and GenAPI [61] have been developed to tackle the problems of mis-assembly and mis-annotation in low-quality draft isolate genomes. They could be very useful alternatives to those older but more popular tools (**Figure 1**) for analyzing MAGs, even although they were not designed for MAGs.

Lastly, the core gene loss of MAGs will naturally affect subsequent analyses that depend on an accurate core gene set. Incorrect core genes will lead to incorrect functional analysis of core genes (**Figure 6**) and phylogenetics analysis (**Figure 7**). COG analysis of core genes helps to understand the survival of species, while the functional study of unique genes are important to understand the strain-specific adaption to environments [38,44]. The accuracy of core/unique genomes is thus the key to the precise interpretation of species survival and environmental adaption. The errors in phylogenetic trees will lead to the misinterpretation of the species evolution and relationship among strains. Clearly, the accuracy of these phylogenetic trees highly depends on the completeness and fragmentation of genomes. The incorrect core genome alignment will further influence the single nucleotide polymorphisms (SNPs) analysis, leading to inaccurate inference of strain relationships. We expect that these challenges cannot be fully addressed by adjusting the core gene threshold and the use of metagenome-aware gene prediction. This demands future development of new pan-genomics tools that will be specifically designed and tested for MAG analysis.

## Conclusions

With the availability of numerous of MAGs (thanks to cheaper DNA sequencing and bioinformatics advances), we have seen an increasing use of MAGs in pan-genomics. Mis-assembly and mis-binning are inevitable with current bioinformatics tools, leading to fragmented, incomplete, and contaminated MAGs. However, no critical assessments of pan-genomics of MAGs have been published. Our paper filled this research gap, and provided a timely and important alert to the microbiome community that pan-genomics of MAGs needs some careful thoughts. We also offered some practical recommendations in the choice of specific pan-genomics tools and parameters to avoid significant accuracy loss. However, ultimate solutions would be the continuous improvement of MAG quality and the development of new pan-genomics tools specifically designed for MAGs.

## Methods

### Literature search of pan-genome analysis with MAGs

We have searched in PubMed and Google Scholar using the keyword query: (“pangenome” OR “pan-genome”) AND (“metagenome-assembled genome” OR “MAGs”) in titles and abstracts. Papers were kept if the following criteria were met: (i) studied organism(s) were prokaryotes, (ii) at least one MAG was used in pan-genome analysis, and (iii) a specific pan-genome analysis tool and specific parameters were described.

### Distributions of fragmentation, completeness, and contamination of real MAGs

A total of 276,349 MAGs of the Unified Human Gastrointestinal Genome (UHGG) collection [38] were used to determine the distribution of contig number, completeness, and contamination in MAGs. Specifically, the MAG summary file in the MGnify FTP site (ftp.ebi.ac.uk/pub/databases/metagenomics/mgnify_genomes/human-gut/v1.0/genomes-all_metadata.tsv) was processed. Histograms and density plots were created using R ggplot2 [62].

### Simulate MAGs from complete genomes

Complete bacteria genomes were downloaded from the NCBI RefSeq database in October 2019 [63]. Only 17 species with at least 100 complete genomes were used to generate simulated MAGs. For each species, 100 complete genomes were randomly selected as the *original dataset*. We simulated MAGs from the complete genomes resembling the distribution of fragmentation, completeness, and contamination observed in UHGG MAGs. For example, to simulate a dataset of 100 MAGs with an average fragment number of 50, a Python script was developed to generate 100 random numbers with a mean = 50 and following an F-distribution. The F-distribution was selected because it fitted the distributions observed in real UHGG MAGs. These 100 random numbers corresponded to the numbers of cuts that were made in the 100 complete genomes to create a *simulated MAG dataset* (100 fragmented genomes). Simulated MAG dataset with incomplete genomes and contaminated genomes were generated in the same manner. The three operations were combined for MAG simulations and depicted in **Figure 2A**.

Fragments from genomes of the same species or genus were added as contamination. This is because contamination in real MAGs is often introduced at the contig binning step, which is based on the fact that DNA fragments of closely related genomes (e.g., of the same species/genus) tend to share more similar nucleotide compositions. The species-level contamination was added from complete genomes within the same species, whereas the genus-level contamination was chosen from genomes of other species within the same genus.

### Combine MAGs and complete genomes to form mixed datasets

MAGs are often combined with complete isolate genomes together for pan-genome analysis. For two representative species, *Escherichia coli* and *Bordetella pertussis*, mixed/combined datasets were created. Four different MAG percentages were used: 10%, 40%, 60%, and 85%. For example, to create a mixed dataset with 40% MAGs, a total of 40 MAGs (40% in 100) would be randomly selected from the simulated MAG dataset (**Figure 2B**).

### Pan-genome analyses of simulated MAGs

Once the input datasets (complete genomes, simulated MAGs, or mixed) were built, proteins were predicted and annotated using Prokka v1.13 [55] with default parameters. Pan-genome analyses were carried out using tools such as Roary v3.13 [15]. Simulated MAGs were generated at different levels of fragmentation, incompleteness, and contamination (**Figure 2**). For example, 5 different levels (50, 100, 200, 300, and 400) of fragmented MAGs were subject to pan-genome analysis. To project the correlation between the number of core gene families and the number of fragments, an exponential model (*y* = *e*^(*ax*+*b*)^) was used, where y is the number of the core gene families in pan-genome and x represents the number of fragments. The predicted fitting curves were plotted using R ggplot2 [62], and the adjusted-R^2^ and P-values were calculated.

### Test the effect of random selection of complete genomes

To select which 100 complete genomes to generate simulated MAGs may affect the analysis results. To assess the variation that may be caused by the selection of different 100 genomes for simulation, 50 random datasets (each with 100 randomly selected genomes) were generated for *Escherichia coli* by going through the same simulation process as depicted in **Figure 2**. For *Bordetella pertussis*, *Staphylococcus aureus*, and *Klebsiella pneumoniae*, 30 random datasets were generated (these species have smaller numbers of complete genomes). These 50 or 30 datasets of the same type (e.g., 100cut fragmentation group) were analyzed separately. The median, mean, and standard deviation for the number of core genes of the 50 or 30 datasets were calculated. The violin plots were created by using R ggplot2 [62] to visualize the variations among datasets.

### Compare three pan-genome analysis tools

Three pan-genome analysis tools, Roary [15], BPGA v1.3 [17], and Anvi’o v7.0 [16], were compared. Two representative species, *E. coli* and *B. pertussis*, were selected for the comparison. To have a fair comparison, all three tools used the same gene prediction files from Prokka v1.13 [55] for the analyzed MAGs, unless stated differently.

Roary was run with the parameters “-i 90 -cd 100 -s -e -n”, where -i defines the sequence identity (SI) threshold in BLAST comparison, and -cd defines the percentage threshold in core gene (CG) definition. These two parameters can be set at different thresholds. In BPGA, USEARCH [64] was selected as the gene clustering tool with the same SI and CG thresholds used in Roary. In Anvi’o, the internal gene prediction was skipped, and a python script was used to feed in the external gene prediction file (Prokka result). There is no SI parameter in Anvi’o, so two equivalent “--mcl-inflation 10 --use-ncbi-blast --minbit 0.8” were used instead.

### Test the use of different SI and CG thresholds in pan-genome analysis

In pan-genome analysis, two parameters are very important: (i) the sequence identity (SI) to define homology (e.g., two genes must be > 90% identical to be clustered into the same homologous gene cluster) and (ii) the core gene (CG) definition threshold (e.g., core genes must be found in 100% of genomes). The simulated MAGs for *E. coli* and *B. pertussis* were used to evaluate the effects of different parameter thresholds on pan-genome analysis results.

Different CG thresholds were compared using Roary, BPGA, and Anvi’o (use the default prodigal): 100%, 99%, 98%, 95%, 92%, and 90%. Different SI thresholds was tested using Roary only: 95%, 90%, 85%, and 80%.

### Test the core gene recovery by lowering CG thresholds

Pan-genome analysis of MAGs will lead to the loss of core genes compared to complete genomes (see Results). Lowering the CG threshold will increase the core gene number, but the recovered CGs may not be the lost ones. To test this, the representative protein sequences of CGs in a simulated MAG dataset were used as the query to search against the representative protein sequences of CGs in the original dataset by using BLASTp.

### Downstream analysis

Two downstream analyses were performed based on the pan-genome results: (i) Clusters of Orthologous Genes (COG) functional analysis and (ii) phylogenetic analysis. The simulated MAG datasets of *E. coli* and *B. pertussis* were used for these two types of analysis.

#### (i) COG analysis

The representative protein sequences of three gene groups (core, accessory and unique genes) were used as the query to search against the COG database [65] using RPS-BLAST with E-value < 1E-5. Genes having multiple non-overlapping domains were assigned to multiple COG functional categories. The number of genes assigned to each COG functional category was calculated. The COG functional enrichment in each gene group was tested by using the one-tailed Binomial test with adjusted P-value >0.05 following our previous papers [66].

#### (ii) Phylogenetic tree comparison

a. Roary provides a phylogeny reconstructed using the presence and absence of accessory genes. It also provides the presence and absence matrix as a dot plot.
b. Roary provides the core gene nucleotide sequence alignment. This alignment was used to construct another phylogeny by using Fasttree v2.1 [67].

To compare two phylogenies and quantify the differences, the normalized Robinson-Foulds (nRF) symmetric distance [68] was used. The nRFs between the tree from the original complete genomes and the tree from simulated MAGs were calculated by using ETE3 toolkit [69].

## Supplementary data

**Table S1**: Provided as an excel document with curated publications

**Figure S1-S9**: Provided as a word document with figure legends

## Declarations

### Ethics approval and consent to participate

Not applicable

### Consent for publication

Not applicable

### Availability of data and materials

All the computer codes and data generated in this study have been made available at GitHub (https://github.com/tli14/PanMAGs).

### Authors’ contributions

T.L. performed all the data analysis. Y.Y. conceived and supervised the study. T.L. and Y.Y. wrote the manuscript. All authors have read and approved the final manuscript.

## Acknowledgements

We thank Dr. Yuzhen Zhou for his suggestion on F statistics. This work was partially completed utilizing the Holland Computing Center of the University of Nebraska–Lincoln, which receives support from the Nebraska Research Initiative.

