## Supplementary material for "Critical assessment of pan-genomics of metagenome-assembled genomes": Figure S1

Yanbin Yin

**Supplementary Figures**

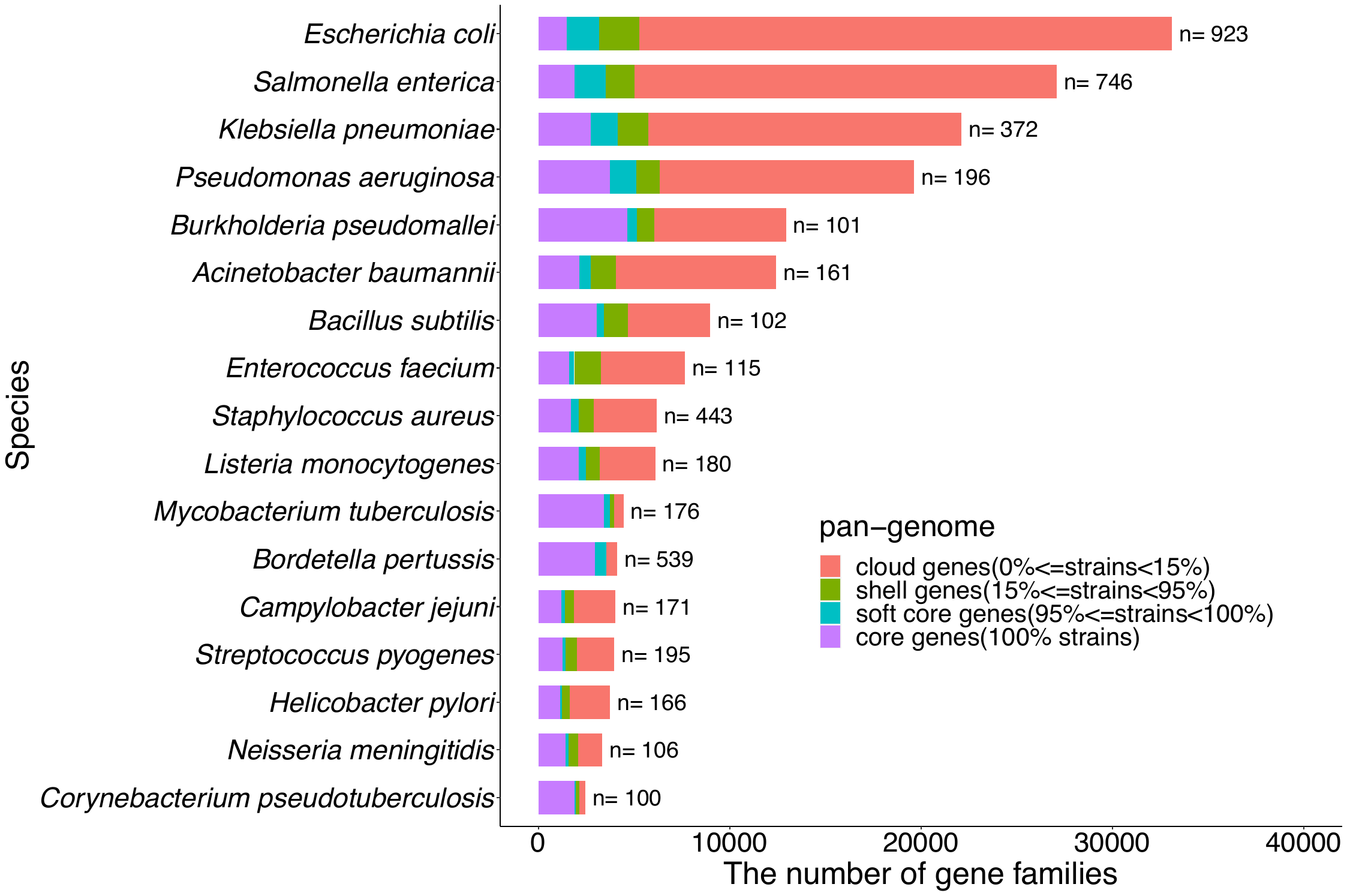

**Figure S1. The Pan-genome composition of 17 species constructed by using Roary.** The n beside each bar represents the number of complete genomes used for building species pan-genome. Roary was run with parameters (-i 90 -cd 100 -s -e -n), which produced four classes of genes shown in different colors. Some terms (e.g., cloud genes and shell genes) are not often seen from other pan-genome analysis tools.

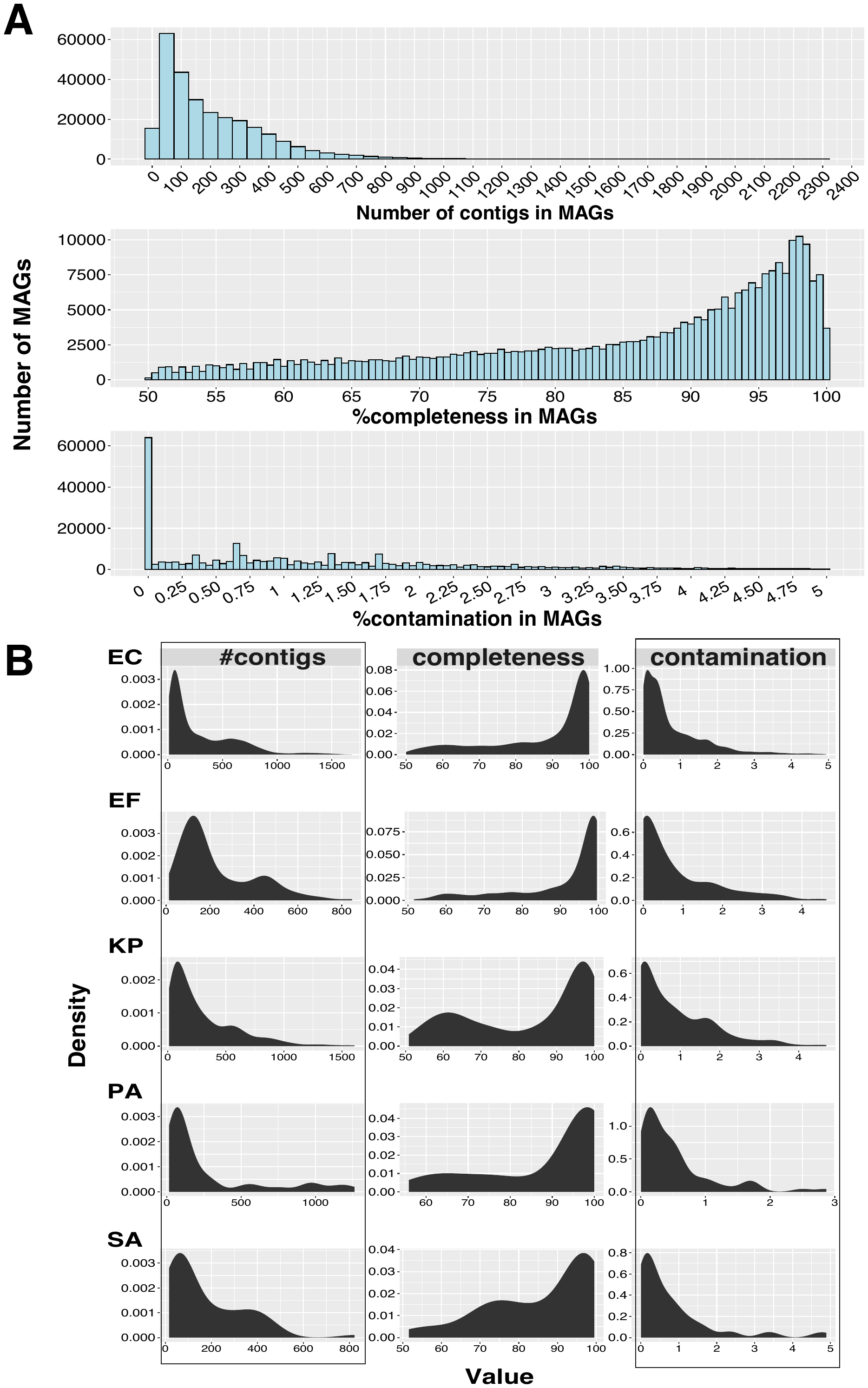

**Figure S2. The statistical analysis of UHGG MAGs.** (**A**): The distribution histogram of contig numbers, completeness percentage, and contamination rates in 276,349 MAGs. (**B**): The density plots for UHGG MAGs of five species: EC for *Escherichia coli* (n=4,391), EF for *Enterococcus faecium* (n=333), KP for *Klebsiella pneumoniae* (n=641), PA for *Pseudomonas aeruginosa* (n=63), and SA for *Staphylococcus aureus* (n=57).

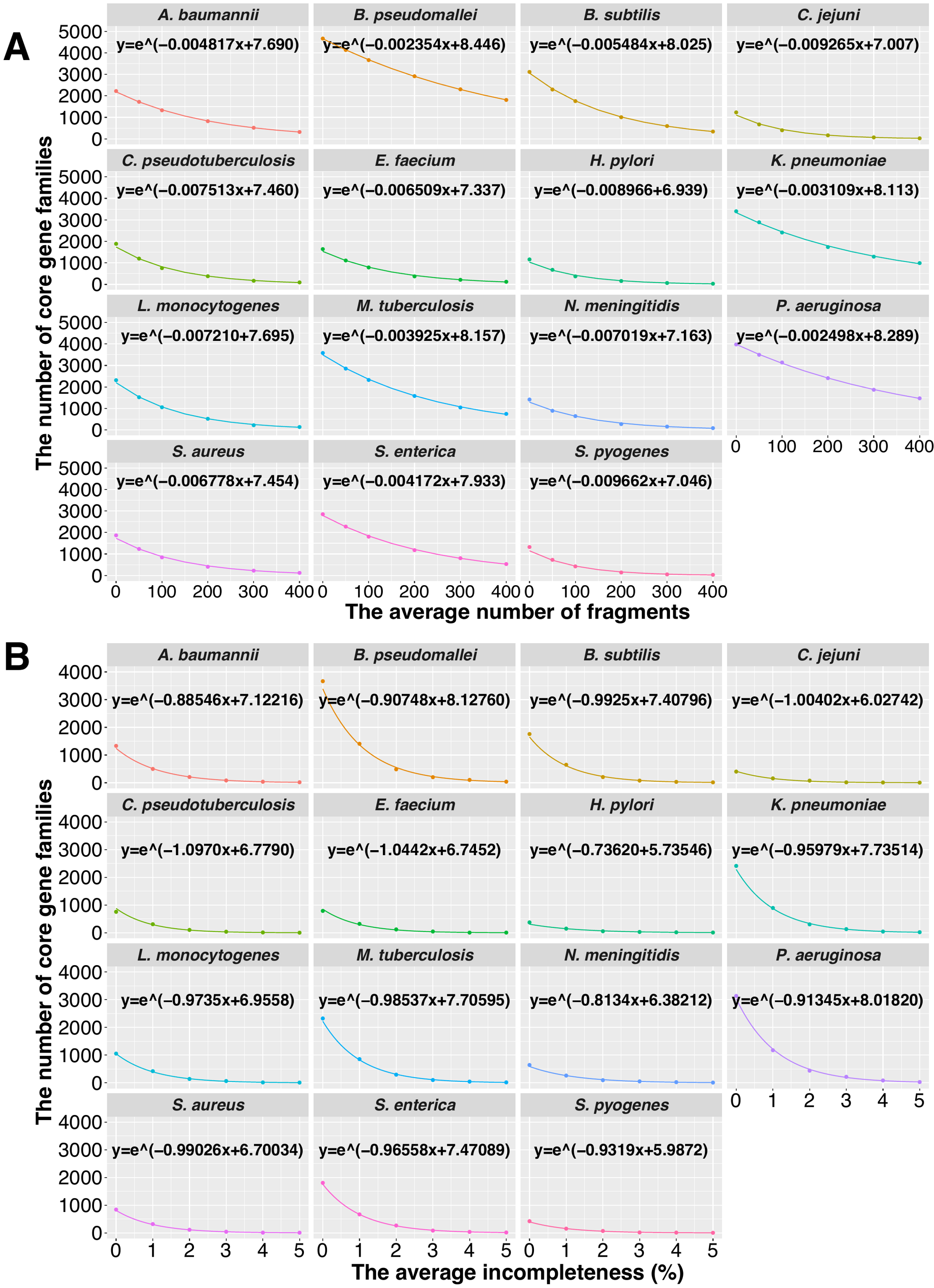

**Figure S3. Fragmentation and incompleteness effect on the number of core gene families**. (**A**): The core genome size continuously decreases as the simulated MAGs become more fragmented. The red curve was predicted using an exponential model for the correlation between the x-axis (the number of fragments) and the y-axis (the core genome size). (**B**): The core genome size decreases more rapidly as the simulated MAGs become less complete.

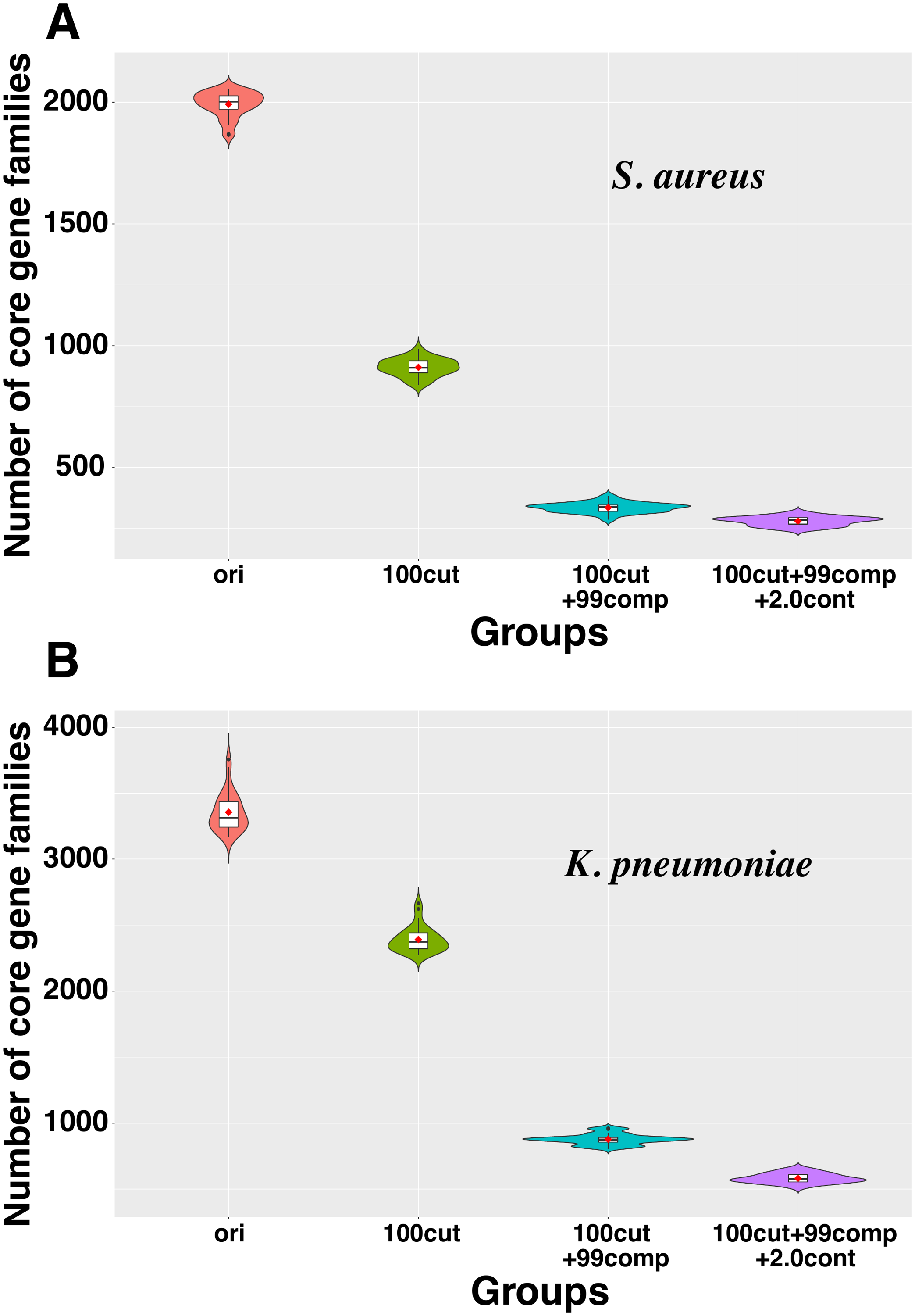

**Figure S4.** Core genome sizes decrease in *S. aureus* and *K. pneumoniae*. (A): The violin plot of the core genome sizes in 30 *S. aureus* original datasets and their corresponding simulated MAG datasets. (B): The violin plot of the core genome sizes in 30 *K. pneumoniae* original datasets and their corresponding simulated MAG datasets. See more details in legend in **Figure 3E and F**.

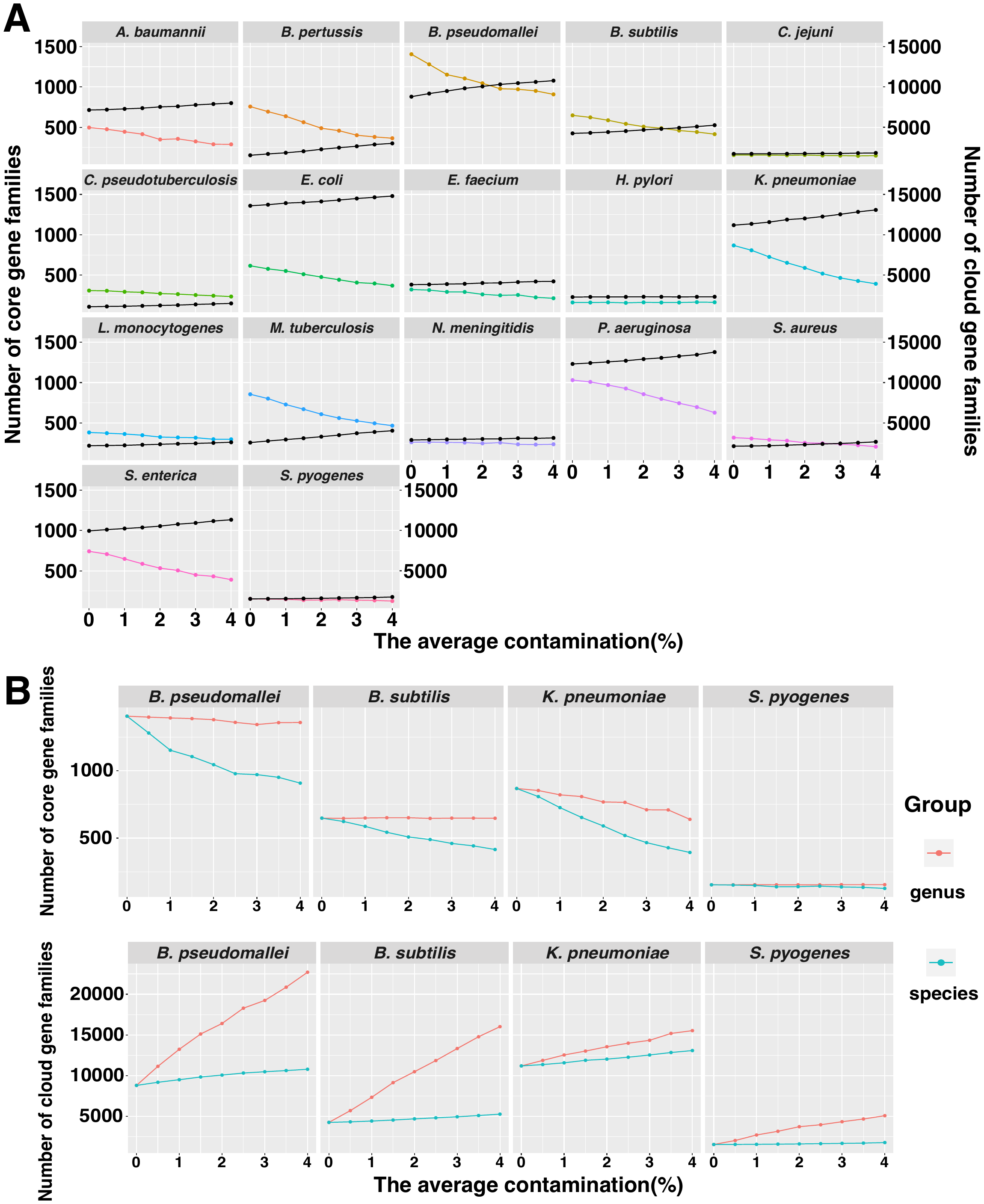

**Figure S5. Contamination effects on the number of core and cloud gene families.** (**A**): The number of core gene families (colored curves, left y-axis) and cloud gene families (black curves, right y-axis) in 100 MAGs with a different average intraspecies contamination rate (%). (**B**): The interspecies and intraspecies contamination comparison. Group labeled as genus represents the interspecies contamination from other species in the same genus, and group labeled as species represents the intraspecies contamination from different strains in the same species.

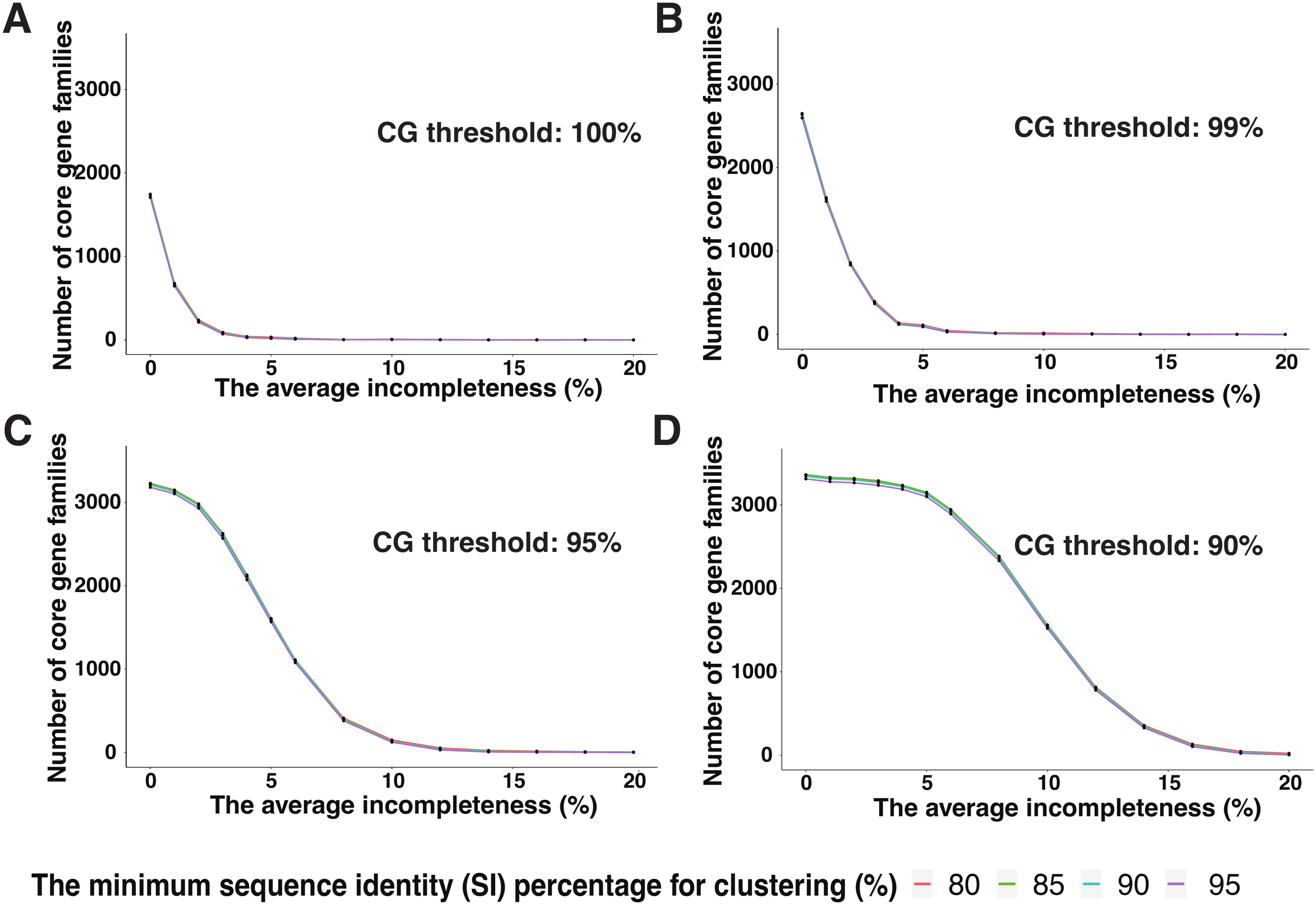

**Figure S6. Different gene sequence identity comparison.** Line plots of the number of core gene families in *E. coli* MAG datasets with different incompleteness percentages. The core gene (CG) threshold used is 100% (**A**), 99% (**B**), 95% (**C**) and 90% (**D**). Different colors represent the minimum identity percentage for gene clustering. In (A) and (B), the four curves almost have no difference and overlap very much.

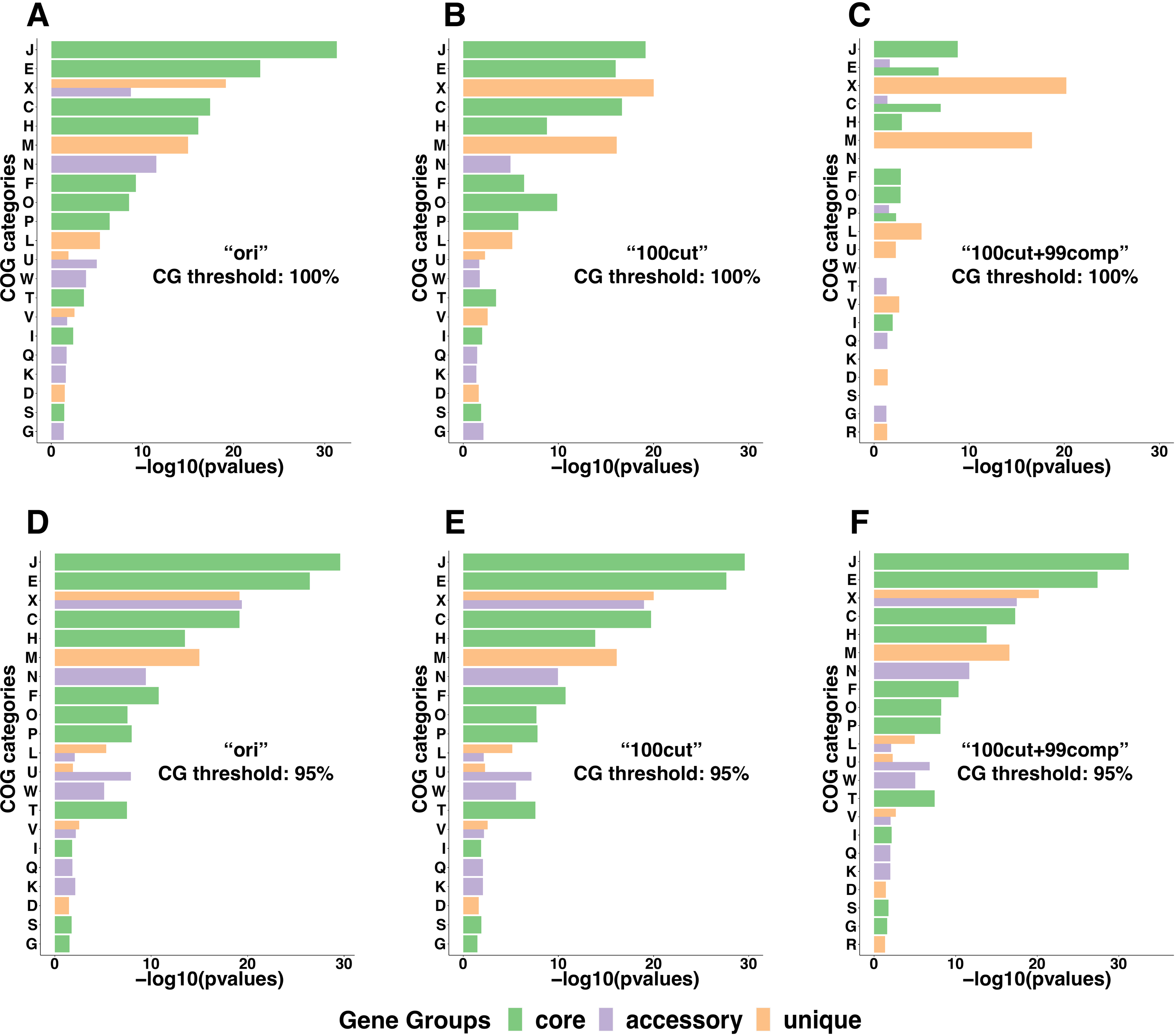

**Figure S7.** **Enrichment of COG functional categories in *E. coli* core, accessory and unique genes compared to the entire pan-genome.** Bar plots show the enrichment of COG categories in each gene group as adjusted P-values (-log10) calculated with binomial tests (adj P-value<0.05). The core gene threshold 100% (**A-C**) and 95% (**D-F**) are used in original datasets and their corresponding simulated MAG datasets. See more details in legend in **Figure 3** and **Figure 6.**

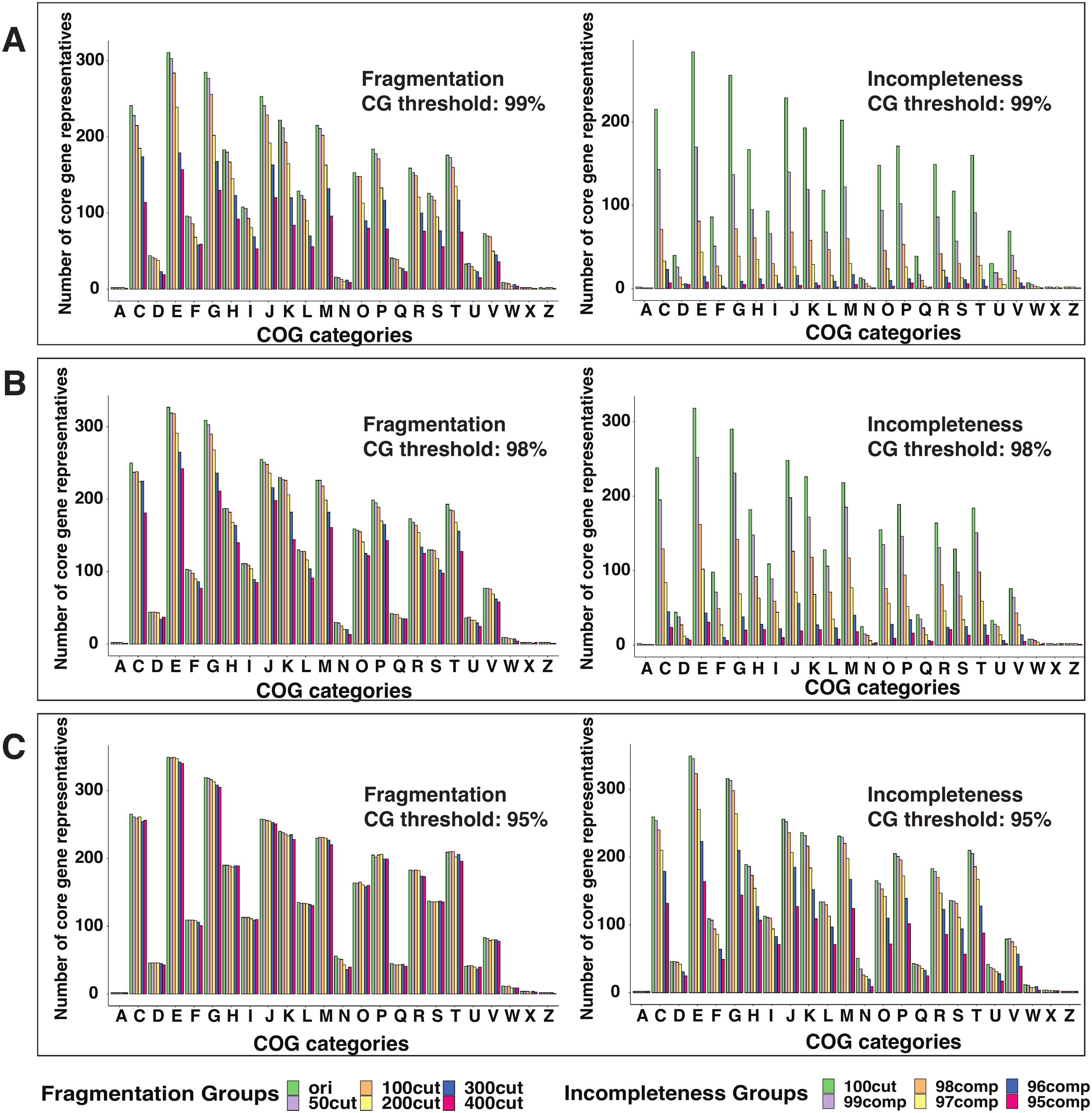

**Figure S8. Core gene thresholds influence COG analysis for *E. coli* core genome.** Bar plots of the number of core gene representatives in *E. coli* datasets with different fragmentation (left) and incompleteness (right) in each COG category. The core gene threshold 99% (**A**), 98% (**B**) and 95% (**C**) are used. See more details in legend in **Figure 6.**

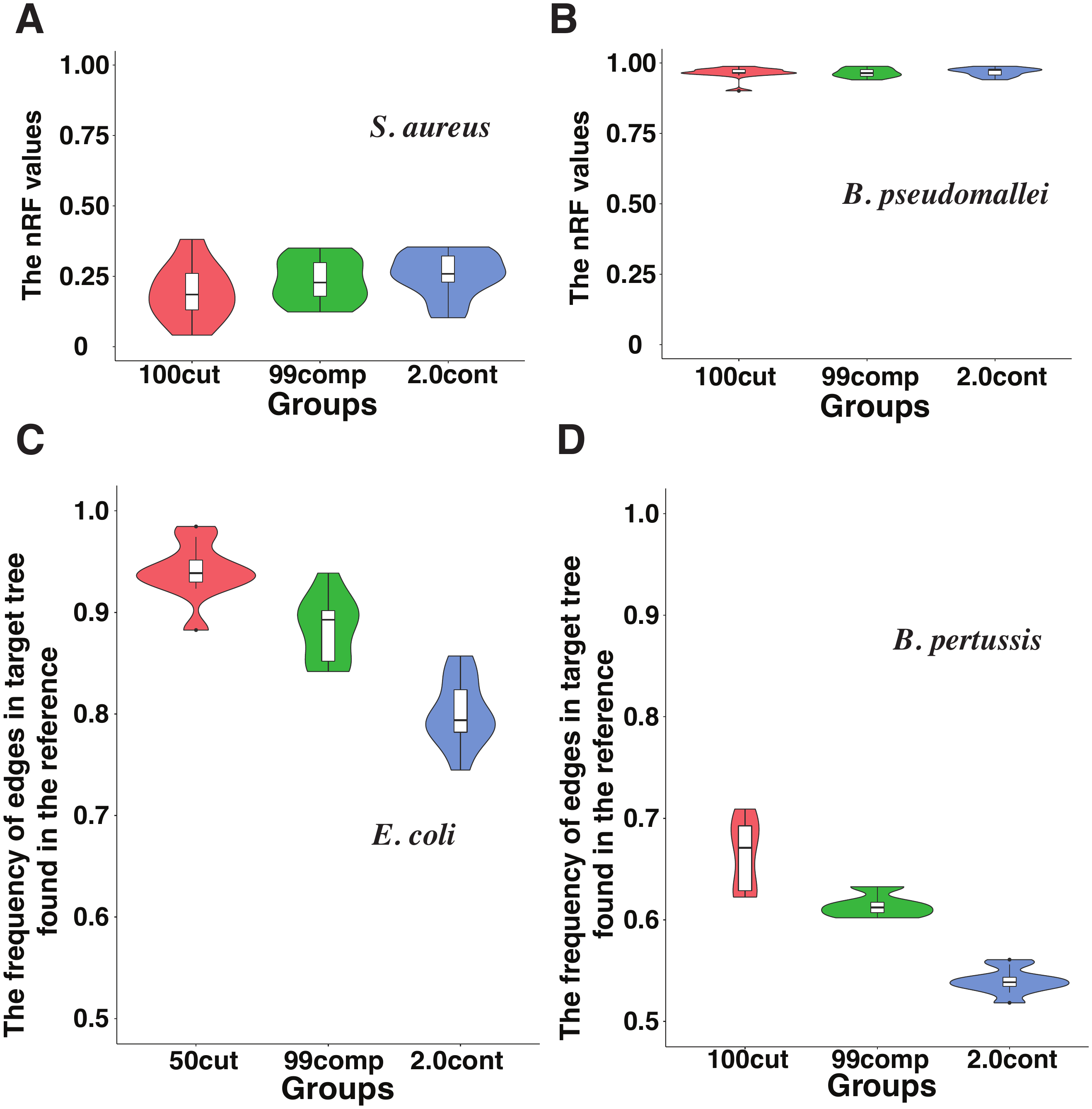

**Figure S9. The effects on phylogenetic trees in different species.** (A) and (B): The violin plots of the nRF distance values between MAG tree and complete genome tree constructed based on gene presence and absence matrix. (**C**) and (**D**): The violin plot of the percentage of tree branches shared between MAG trees and the complete genome tree constructed based on core genome alignment.
